# Mimicking the night shift: Hypothalamic-pituitary-adrenal (HPA) axis regulation, energy homeostasis, and oxidative stress in a model diurnal songbird

**DOI:** 10.1101/2025.06.09.656008

**Authors:** Kevin Pham, Victoria M. Coutts, Alexander J. Hoffman, Haruka Wada

## Abstract

Everchanging socioeconomic demands, coupled with modern day technological advancements have lifted constraints on human activity patterns, enabling us to take advantage of the 24-h period. However, varying patterns of light at night and sleep/wake may come at the cost of disrupting endogenous rhythms that govern biological processes and maintain homeostasis. To this end, we exposed zebra finches (*Taeniopygia guttata castanotis*), a diurnal songbird, to a consistent light/dark cycle of 12L:12D or alternating lighting patterns of 12D:12L and 12L:12D every three days for 66 days to mimic night shift work conditions. Surprisingly, we found that corticosterone profiles were inverted in birds exposed to simulated night shift work, with a slight trend indicating the inversion of melatonin profiles. Moreover, they had poor HPA axis reactivity in response to a standardized capture and restraint challenge and a significant reduction in corticosterone levels at ZT0 after simulated night shift work. Yet, we did not detect changes in body mass, subcutaneous fat deposits, or 4-hydroxynonaneal damage in the brain or liver. Our study underscores the need to capitalize on unique model organisms to evaluate consequences and resilience of simulated night shift work to inform public health decisions and the role of circadian plasticity.

## Introduction

Loss of adequate sleep quality and duration has become one of the most prominent drivers of circadian rhythm disruption in today’s industrialized world. Sleep disturbance results from a variety of sources, including social jetlag or prolonged exposure to bright light prior to sleep onset, or shift work during the dark phase (Guindon et al., 2024; James et al., 2017). Exposure to distinct light spectra can differentially stimulate photoreceptors/pigments in the retina, pineal gland, or suprachiasmatic nucleus (SCN), causing signaling cascades that activate or inhibit multiple endocrine axes (Ouyang et al., 2018). Consequently, conflicting patterns of light at night can result in temporal mismatch between master and peripheral oscillators that phase shift core clock genes, alter gene expression profiles across the transcriptome, and disrupt hormone regulation (Husse et al., 2017; Kervezee et al., 2019).

There are multiple scenarios where an individual is exposed to light at unnatural times or experiences wakefulness during the dark phase. One scenario is through shift work, defined as any shift outside 7am-6pm (Caruso, 2014). Consistent or rotating night shifts encompass hours outside the common day shifts, at various patterns, resulting in alteration between a certain number of days on night shift then days off shift or to a different type of shift (Pilcher et al., 2000). During this time, tasks and recreational activities can greatly vary between individuals that consist of different psychosocial pressures that result in sleep loss, among other mental strains such as anxiety or irritation (Costa, 2003; Pilcher et al., 2000).

Night shift workers span across multiple sectors such as healthcare or transportation. In fact, epidemiological studies in health care workers show increased signs of pathologies associated with insulin resistance or obesity that predispose them to developing metabolic syndromes (Caruso, 2014; James et al., 2017). Yet, one critical gap in our knowledge stems from the fact that there are inconsistencies in cross-sectional and observational epidemiological studies that link night shift work (among other things) to disease outcomes due to confounding variables and lack of statistical power (Asiamah et al., 2021; Hansen and Pedersen, 2025; Van et al., 2021). Because shift work spans across multiple sectors, interpersonal social interactions or work intensity can magnify the burden of night shift work or mask the effects.

Perhaps one of the main limitations regarding manipulative experimental studies regarding shift work, is that it primary focuses on rodent models (Opperhuizen et al., 2015). Although these studies have proven useful in linking similar and clear patterns regarding glucose intolerance (*e.g.,* hyperglycemia) and melatonin suppression between mammalian species, there are some studies that show differential effects between mice and rodents under the same circadian rhythm disruption paradigm (Summarized in Opperhuizen et al., 2015). Additionally, rodents have opposing phase relationships compared to humans such as glucocorticoid secretion, activity on/offset, and sleep/wake cycles. Therefore, translating findings from nocturnal rodents to diurnal humans may not be fully representative and should be done with caution (Mendoza, 2021; Opperhuizen et al., 2015).

Nevertheless, longitudinal, cross-sectional, or cohort based epidemiological studies have provided a solid foundation of evidence clarifying that night shift work is indeed a detrimental stressor (Guindon et al., 2024; James et al., 2017; Walker et al., 2020). Such associations with the development of diseases has prompted researchers and the medical community to redefine night shift work as a carcinogen, given its link to the development of multiple cancers even after controlling for biases and confounding factors (World Health Organization monographs, 2019; Moon et al., 2024). However, there are likely alternative physiological outcomes and patterns in response to night shift work based on sex, chronotype, differences in sensitivity, and temporal niche, given that manipulative studies in diurnal animals are lacking (but see Mendoza, 2021).

Birds provide a unique opportunity to explore the effects of shift work and circadian misalignment. Songbirds have phase relationships that closely mimic human sleep/wake cycles and similar rhythms in glucocorticoid and melatonin secretion (Bahammam, 2025; Cassone, 2015, 2014). In contrast to mammals of similar body size, birds have a substantially higher resting metabolic rates, which reflect almost a 2-3 fold difference in fasted glucose levels and the capacity to mount glucose responses that exceed point-of-care device detection limits (600 mg/dL) (personal observations) (Braun and Sweazea, 2008; Sweazea, 2022). Yet, these extreme levels of glycemia show little to no association with oxidative stress or the glycation of biomolecules in birds (Brun et al., 2022; Sweazea, 2022). This is in direct opposition to rodent and human models, that are unable to sustain hyperglycemic conditions, known to be a driver in the development chronic health diseases, longevity, and damage (Sweazea, 2022). Thus, a bird’s natural evolutionary resistance to hyperglycemia and cellular damage provides an interesting avenue to investigate the mechanisms that may underlie their resilience to metabolic disruption in response to simulated night shift work, with the perspective of the evolutionary pressures that have shaped their life-history.

Zebra finches (*Taeniopygia guttata castanotis)* are an arid-adapted, diurnal songbird native to Australia. For decades, they have been the prime candidate model to evaluate endocrine axes, such as the hypothalamic-pituitary-adrenal (HPA) axis, which regulates physiological stress responses that mediate plastic responses to environmental variation, among other endocrine axes that underlie the neurobiology of learning and cognition (Perfito, 2010; Spierings and Cate, 2016). Unlike other diurnal birds that show flexibility in their nighttime activity (and possibly their physiology), zebra finches are strictly diurnal, showing little to no activity during the dark phase (Wang et al., 2012). Zebra finches are also opportunistic songbirds, meaning that photoperiodism is generally not the ultimate cue to time crucial life history events such as reproduction (Perfito et al., 2008). Thus, they provide the perfect opportunity to disentangle how shifting patterns of light at light underpin specific neuroendocrine pathways and their downstream consequences, without activating or confounding other pathways sensitive to photoperiodism and associated with life-history variation (gonadal stimulation).

To fill critical gaps in both rodent and human night shift literature, we leveraged the unique biology of the zebra finch to investigate how simulated night shift work altered regulation of two key hormones: glucocorticoids and melatonin. Glucocorticoids and melatonin are both under circadian control, and reflect circadian alignment, albeit downstream from the master clock. The zenith of glucocorticoids prior to activity onset or the early morning primes the body for wakefulness by increasing blood pressure, alertness, and heart rate, while simultaneously increasing appetite and promoting catabolism. Conversely, the nadir of glucocorticoids is thought to depress the body for the predicted resting phase by decreasing blood pressure and slowing down heart rate, while suppressing appetite and promoting anabolism (Oster et al., 2017; Rao and Androulakis, 2019). While melatonin is critical in biological timekeeping, it also acts as an antioxidant (Bonnefont-Rousselot and Collin, 2010). The melatonin antioxidant cascade integrates this hormone as well as its metabolites to reduce the superoxide anion (O2-), a normal byproduct of cellular metabolism, prior to or after chemical reactions with other reactive species such as hydroxyl and peroxyl groups (Tan et al., 2007). Thus, in combination with other endogenous antioxidants (e.g., superoxide dismutase) melatonin serves as a multifunctional hormone to prevent oxidative stress and accumulation of damage.

In this study, we exposed 24 female adult zebra finches to alternating light cycles of 12D:12L and 12L:12D to mimic rotating night shift work conditions for a total of 66 days, with 3 days on the simulated night shift protocol and 3 days off (while other 24 females remained on 12L:12D; N=48). We hypothesized that simulated night shift work would alter regulation of the hypothalamic-pituitary-adrenal (HPA) axis and melatonin rhythms, resulting in downstream physiological consequences on energy metabolism, morphometrics, and oxidative stress.

We predicted that chronic exposure to our treatment paradigm would blunt the adrenocortical response and reduce their ability to facilitate negative feedback, resulting in poor efficiency in the breakdown of energy substrates (glucose and β-hydroxybutyrate). Furthermore, we predicted that the resulting disruption in energy homeostasis would lead to a rise in body mass and subcutaneous fat deposits. We also predicted that simulated night shift work would suppress the diurnal peak of glucocorticoids and the nocturnal peak of melatonin. Although we expected a suppression of these hormone peaks, we predicted that treatment birds would sustain higher levels of glucocorticoids and lower levels of melatonin compared to controls, therefore increasing the prevalence of oxidative damage in the liver and brain.

## Methods and materials

### Animal husbandry

Adult female zebra finches (N = 48) (all <1 year old) in this study originated from a captive breeding colony at Auburn University with known hatch date and history as well as from a local breeder located in Georgia, USA (∼2 hours away from AU) and have never previously experienced light at night. Zebra finches were housed in in tower cages (38.10 cm width × 45.72 cm depth × 45.72 cm height) and maintained under a 14L:10D photoperiod at Avian Research Laboratory II at Auburn University. Average temperature and humidity levels ranged between 21.92°C (± 2.18 SD) and 58% (± 22 SD), respectively. A mixture of finch seeds (Kaytee supreme) and water was provided *ad libitum*.

Finches from both origins (20 birds from our breeding colony and 28 birds from the local breeder) were weight-matched and randomly distributed into two experimental groups. Animals were then transported from Avian Research Laboratory II to the Biological Research Facility on AU’s campus (<1.2 miles) and housed there for the remainder of the experiment.

Once at the Biological Research Facility, zebra finches were further distributed and housed across four identical, temperature and humidity controlled animal housing rooms (2 rooms/treatment; n = 12/room). To account for potential differences due to Origin, birds from both sources were equally distributed across treatment groups and the animal housing room, with five AU birds and seven from the local breeder in each. Birds were singly housed in Prevue Pet Products black wire cages (24 in. L x 16 in. W x 16 in. H with 3/8 in. wire spacing) separated by a cage divider and situated on secured metal racks. Animals were within sight of one another, and we followed the same animal husbandry regimen that was conducted at Aviary Research Laboratory II.

Birds were able to acclimatize to their new environment and photoperiod set at 12L:12D (lights on 6am; lights off at 6pm) for at least three weeks before pretreatment samples were collected. The lighting source provided were PLT Solutions 17-Watt 5000K white LED lights which consists of shorter wavelengths of light known to disrupt glucocorticoid levels at low-level light intensities and activate deep brain photoreceptors in avian species (Alaasam et al., 2018). Control birds experienced an average of 34.83 lux/ft^2^ (± 19.17 SD), while treatment birds experienced an average of 36.51 lux/ft^2^ (± 15.29 SD), measured using Hobologgers (HOBO onset product # UA-002-64 and U12-006). Low ambient white noise (<10 decibels) was played throughout the entirety of the experiment to eliminate unwanted outside noise. All animal handling, training, and care procedures were approved by the Institutional Animal Care and Use Committee (IACUC) protocol #2023-5154 at Auburn University.

### Experimental design and sampling scheme

All birds were acclimatized to a 12L:12D light cycle with lights turning on at 6am (ZT0) and lights turning off at 6pm (ZT12). To simulate night shift work, the treatment group (n = 24) switched between a 12D:12L and 12L:12D light cycle every three days when experimental treatment began (**Fig. 1)**. This resulted in treatment birds experiencing “dark hours” starting at 6am and “light hours” starting at 6pm. After three days of the night shift protocol, treatment birds switched back to a 12L:12D light cycle to mimic “off shift” conditions, in which they experienced the same light cycle as controls and were previously acclimated to, and this pattern persisted for the duration for the experiment (66 days). Animal husbandry was completed only when animals experienced light hours. Moreover, sampling for our variables of interest only occurred when all animals experienced the same light cycle.

**Figure 1.**
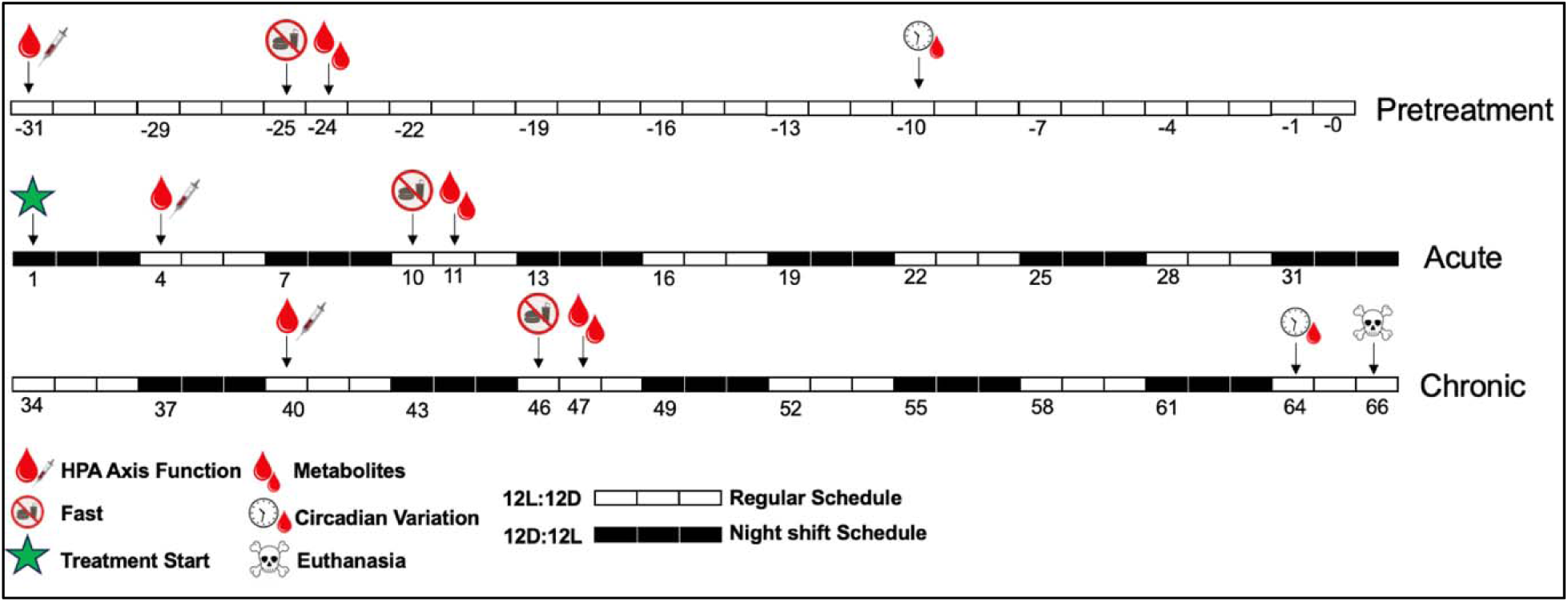
Experimental schematic of pretreatment sample collection. Each singular white bar represents one day on a regular light/dark schedule (lights on at 6am with lights off at 6pm). Numbers with a negative sign indicate days prior to treatment start (pretreatment). Blood icon with syringe on day −31 indicates when pretreatment HPA axis dynamics was tested (baseline, post-restraint, and post-dexamethasone corticosterone levels). On Day −25, birds were fasted 1 hour prior to lights off at 6pm, which is indicated by a crossed out burger and shake icon and bled the following morning on day −24 (one week later from HPA axis dynamics measures) indicated by double blood icons. Circadian variation in circulating hormones were measured on day −10 (two weeks later), indicated by the clock and blood icon. The green star indicates treatment start, in which birds experienced their first night shift condition by experiencing darkness from 6am-6pm (ZT0-ZT12pm). Then, light hours began from 6pm-6am (ZT12-ZT24). From here, three solid white bars indicate three days on a regular light/dark schedule (12L:12D) and three solid dark bars indicating three days on a night shift schedule (12D:12L). This schedule persisted for a total of 66 days and light schedules switched every three days. Euthanasia and tissue collection is indicated by the skull and crossbones.

Per IACUC regulations (Protocol #2024-5154), to not exceed 140 µL of blood sample collection every 2 weeks (*i.e.,* 1% of an average 14g zebra finch’s body mass), blood sampling for different metrics occurred on different days. We measured HPA axis dynamics after 3 and 21 night shifts, which were on days 4 and 40 of the experiment, respectively, (**Fig. 1)**. HPA axis dynamics included baseline, post-restraint, and post-dexamethasone regulation of circulating corticosterone (the main avian glucocorticoid and hereby referred to as Cort) levels. Blood samples (50µL) were collected within 3 min of disturbance via the brachial vein using a 26 gauge needle. Birds were subsequently restrained in a brown opaque bag as a standardized inducer of the glucocorticoid stress response. After 30 min, a post-restraint blood sample (30 µL) was collected from finches. Immediately after the post-restraint blood sample was taken, feathers covering the left pectoral muscle were gently pushed aside and the area was sterilized with a 70% ethanol-soaked cotton ball. After allowing the ethanol to evaporate (<15 seconds), dexamethasone (a synthetic glucocorticoid) was administered to individuals intramuscularly using a 0.3 mm insulin syringe, prepared at a dosage of 1000 µg/kg of body weight (Jimeno et al., 2018; Kriengwatana et al., 2014). Finches were returned to their home cages to initiate negative feedback. After 30 min, we obtained a final blood sample (50 µL) to evaluate glucocorticoid negative feedback efficiency. All blood samples were spun at 14,800 *x g* for 10 min to separate plasma and red blood cells, snap frozen in liquid nitrogen, and subsequently stored in a −80°C freezer until analysis.

Fasted levels of blood metabolites (glucose and β-hydroxybutyrate) were measured after 6 and 24 night shifts on the morning of experimental days 11 and 47, respectively, following a brief fast from the night prior (**Fig. 1)**.

Intraindividual variation in circulating hormone levels were measured prior to experimental manipulation and after 64 days of chronic simulated night shift work. Blood samples were collected from different individuals (n= 4/treatment) within 3 min of disturbance across six collection timepoints every four hours (ZT0, ZT4, ZT8, ZT12, ZT16, ZT20). ZT0 corresponds to 6:00am. Between ZT12-ZT20, finches were bled in under red light in total darkness to prevent melatonin suppression. After blood collection, samples were immediately spun down at 14,800 *x g* for 10 min to separate plasma and red blood cells, snap frozen in liquid nitrogen, and subsequently stored in a −80°C freezer. Birds were euthanized via isoflurane two days later on day 66 and we collected the right hemisphere of the brain and liver all within ten minutes of death.

Due to unknown reasons, two individuals from the control group died during the experiment. Therefore, the limited data collected from these birds were removed from all analyses.

### Morphometrics

Body mass (to the nearest 0.01 g) and a visual score (scale of 0-5) of the subcutaneous fat deposits over the tracheal pit (furculum fat) was obtained seven days prior to HPA axis dynamics sampling days. This ensured as accurate as possible a weight measurement to adjust injection volumes for correct dosage of dexamethasone across all birds, while also minimizing disturbance too close to the sampling timepoint. Body mass measurements were obtained using a scientific scale and fat was scored by a single observer (CITE). To account for variance in body size, we measured the left and right tarsus length to the nearest 0.01 mm using calipers by a single observer.

### HPA axis dynamics

We measured HPA dynamics across three collection timepoints (Pretreatment, 3, and 21 night shifts) using Arbor Assay Corticosterone Assay kit (Ann Arbor, MI cat # K014-H5). Following validations in our lab, we diluted plasma samples 1:10 with 8% disassociation reagent and ran samples in duplicate such that each unique individual measured across all collection timepoints were represented on a singular plate. A nine-point standard curve was generated in triplicate from 10,000 pg/mL to 39.06 pg/mL. A known concentration of Cort standard was prepared at 625 pg/mL and aliquoted across all plates to assess interplate variation. Due to unexpectedly low levels of Cort, all baseline samples (n = 136) fell off the standard curve and were undetectable; therefore, we removed them from the further analysis. Negative feedback efficiency was calculated as the relative reduction or percent change from restraint-induced Cort to post-dexamethasone (Post-dex) Cort using the following equation below (Lattin et al., 2020). Positive values indicate that individuals suppressed Cort, with 100% indicating complete inhibition, while negative values indicate individuals secreted Cort, even after the dexamethasone injection. A total of 15 plates were ran across two days. Intraplate variation was 7.1% while interplate variation was 9.1%.

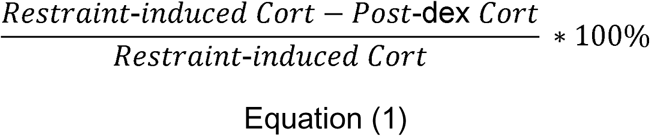

### Metabolites

Following a brief fast from the night prior, blood was collected within 3 min of disturbance the subjective morning to measure circulating metabolites. Blood glucose (mg/dL) and β-hydroxybutyrate (mmol/L) levels were measured in duplicate using the ReliOn prime blood glucose monitoring device (Walmart Apollo LLC Bentonville, AR) validated in our lab (Pham et al., 2025) and the PrecisionXtra ketone monitoring device (Abbott; product #9881465), respectively.

To validate the PrecisionXtra ketone monitoring device for use in zebra finches, we compared β-hydroxybutyrate levels measured between the point-of-care device and a bench assay to ensure values collected by the meter were reflective of values from the bench assay (supplementary materials; **Fig S1**).

### 24-h circulating hormone level

Because we expected low levels of circulating corticosterone throughout a 24-h period, we utilized an alcohol extraction method to extract corticosterone from plasma following (Parks et al., 2023) with slight modifications. Rather than 5 μL of starting plasma, we used 10 μL to increase our precision and ability to detect low levels of corticosterone with 40 μL of assay buffer provided by the Arbor Assay Corticosterone Assay kit (Ann Arbor, MI cat # K014-H5). Afterwards, 250 μL of ethyl-acetate was added to tube and was vortexed for 30 s at 1800 RPM. Samples were allowed to incubate for 5 mins at room temperature to allow for phase separation. Samples were then placed in the −80°C freezer for 10 minutes. After 10 min, samples were taken out, allowed to thaw, and the alcohol layer was collected. We repeated this process two more times for a total of 600-700 μL of the alcohol layer collected. To calculate extraction efficiency, a plasma pool from non-experimental bird samples was stripped of hormone and spiked with a corticosterone standard (100,000 pg/mL) to a known concentration of 10,000 pg/mL. We followed the same extraction method for this spiked sample. All samples were allowed to fully evaporate overnight under a fume hood.

After evaporation, tubes were reconstituted with 125 μL of assay buffer and placed in the −20°C freezer until assayed two days later using Arbor Assay Corticosterone kit (Ann Arbor, MI cat # K014-H5). A total of four plates were ran, with each unique individual measured across the two collections timepoints represented on a singular plate. Because we recovered 96% of the corticosterone from the spiked sample, we adjusted raw values based on 100% extraction efficiency to be comparable to other studies measuring 24-h corticosterone, and thus, report data based on 100% efficiency. Intraplate variation was 1.3% while interplate variation was 2.6%.

To measure 24-h melatonin, we used a Melatonin ELISA Kit (OKEH02566) previously validated in zebra finches (Mishra et al., 2019). We diluted plasma samples 1:1 with the sample dilutant provided by the kit and ran the assay according to the manufacturer instructions. A 7-point standard curve was generated in triplicate from 1,000 pg/mL to 15.63 pg/mL. Due to plasma limitations, we ran our samples as singlets, however, our values are reflective of previously published literature using the same kit in zebra finches across similar ZT collection timepoints (Alaasam et al., 2021). Intraplate variation was 9.8%.

### Immunoblots

To prepare samples for immunoblotting, we utilized the right lobe of the liver and the right hemisphere of the brain, after removing the cerebellum, optic tectum, and a small portion of the frontal lobe, such that the majority of the brain tissue utilized was the diencephalon. Liver tissue was prepared in the same manner as previously described (Pham et al., *in press*; Hoffman et al., 2024), but due to low protein abundance in the brain, we standardized protein concentration to 0.5ug/ul across all brain samples. Primary antibodies included an anti-rabbit polyclonal 4-hydroxynonaneal antibody (Abcam 1:1000) and anti-rabbit polyclonal SOD1 antibody (GeneTex1:500) previously. The secondary antibody was an anti-rabbit horse radish peroxidase antibody prepared at 1:2000 for 4-hydroxynonaneal and 1:1000 for SOD 1. Protein bands were visualized via chemiluminescence using ECL prime and images were captured using LAS 4000. We used Image Analysis (Bio-Rad) to perform densitometry. We used ponceau staining to normalize values to total protein.

### Statistical analysis

All statistical analyses were run in R (Version 4.5.0). Statistical significance was determined using an alpha value of *P* ≤ 0.05. Global Linear mixed effects models using the “lme4” and “lmertest” package were generated with Treatment (two level factor) and Timepoint (three level factor) treated as fixed effects as well an interaction term between the two variables (Treatment * Timepoint). Fixed covariates included body mass and tarsus length to control for variation in body size and fluctuations in body mass. For β-hydroxybutyrate and glucose response variables, we also included a fixed covariate of order to control for the sequence in which we entered animal housing rooms during the metabolite measurement days. A fixed factor of origin was also included in all global models in order to determine whether patterns of our response variables were associated with the source population of our birds. To evaluate changes in fat score, we used the “ordinal” package in R (CITE), as each incremental value in the score represents a change in the amount of subcutaneous fat. We included individual Bird ID as a random intercept in all models that had response variables that were sampled across multiple timepoints to account for repeated measures. For immunoblots, gel was included as a random effect. Protein abundance of SOD and 4-HNE were log-transformed to meet assumptions of the linear model. If a significant interaction term was detected on any of the response variables, Tukey’s post-hoc test followed using the “emmeans” package in R for pair-wise comparisons. Non-significant interactions and covariates were removed from global models, and thus, the final model with results reported is the simplest model.

We used the “GLMMcosinor” package in R to fit models on 24-h hormone variables. This package combines the power of generalized linear models to fit multiple types of data distributions in addition to rhythmic data that often violate the assumptions of linearity and normality. Furthermore, because we had birds measured at the same ZT time before and after exposure to treatment we examined within-individual differences using linear mixed effects models at specific ZT times. In regard to the cosinor analysis, the response variables included circulating plasma corticosterone and melatonin collected across six sampling timepoints, and dependent variables ZT time, Treatment, and a family = Gamma or Gaussian distribution for corticosterone and melatonin, respectively. Shapiro-Wilks test indicated non-normality of 24-h corticosterone and we chose a gamma distribution based on the best fit theoretical distribution using simulated data (n = 1000 iterations) that was generated from raw data via the “thefitdistrplus” package in R. Residuals of melatonin were normally distributed, and thus a Gaussian distribution was sufficient. We tested for group differences based on treatment on the acrophase (the peak of the hormone), amplitude (peak hormone level subtracted from average hormone levels), and mesor (the average rhythm of hormone). The acrophase of corticosterone and melatonin was calculated by extracting the beta estimate measured in radians divided by 2rr multiplied by the period (24) following the below equation.

In regard to the linear mixed effects models, we had specific *a priori* predictions that individuals under night shift work would have a suppression in their corticosterone peak that coincides with ZT0, the predicted zenith for corticosterone release and at ZT12, the predicted nadir. Thus, we ran our models by subsetting data by specific ZT times. The response variable included 24-h corticosterone with the interaction of treatment and collection timepoint (pretreatment/exposure; 2 level factor). Bird ID was included as a random effect to account for non-independence and repeated measures. Because we had data points missing for pretreatment 24-h melatonin profiles, we lacked statistical power to investigate this hormone in a similar manner.

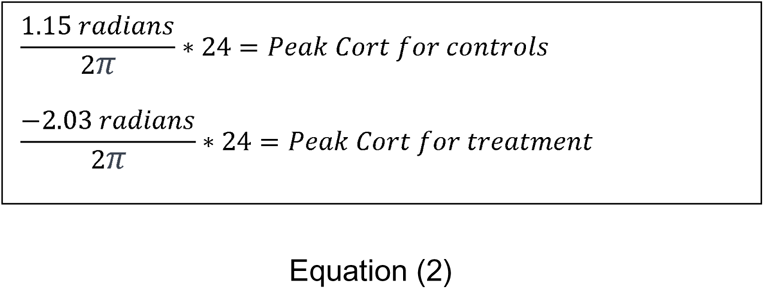

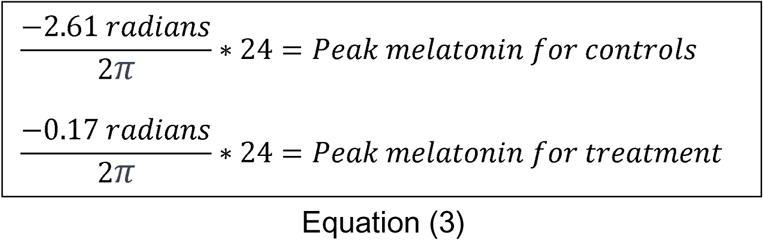

## Results

### Circadian variation in hormone levels

Linear mixed effects model indicated a significant interaction between collection timepoint and treatment (F_1,5_ = 6.55; *P* = 0.05) on corticosterone levels specifically collected at ZT0. Post-hoc analysis found that birds under chronic night shift work had an average suppression of −1.93 ng/mL in their corticosterone levels between the two collection timepoints (Tukey’s Post hoc: β = −1.93; 95% CI = −4.21 to −0.65; *t* = −2.69; *P =* 0.04; **Fig. 2A**). We found that 87% of the overall variance in 24-h corticosterone levels at ZT0 was due to within-individual variation, while only 13% was due to between-individual differences.

**Figure 2.**
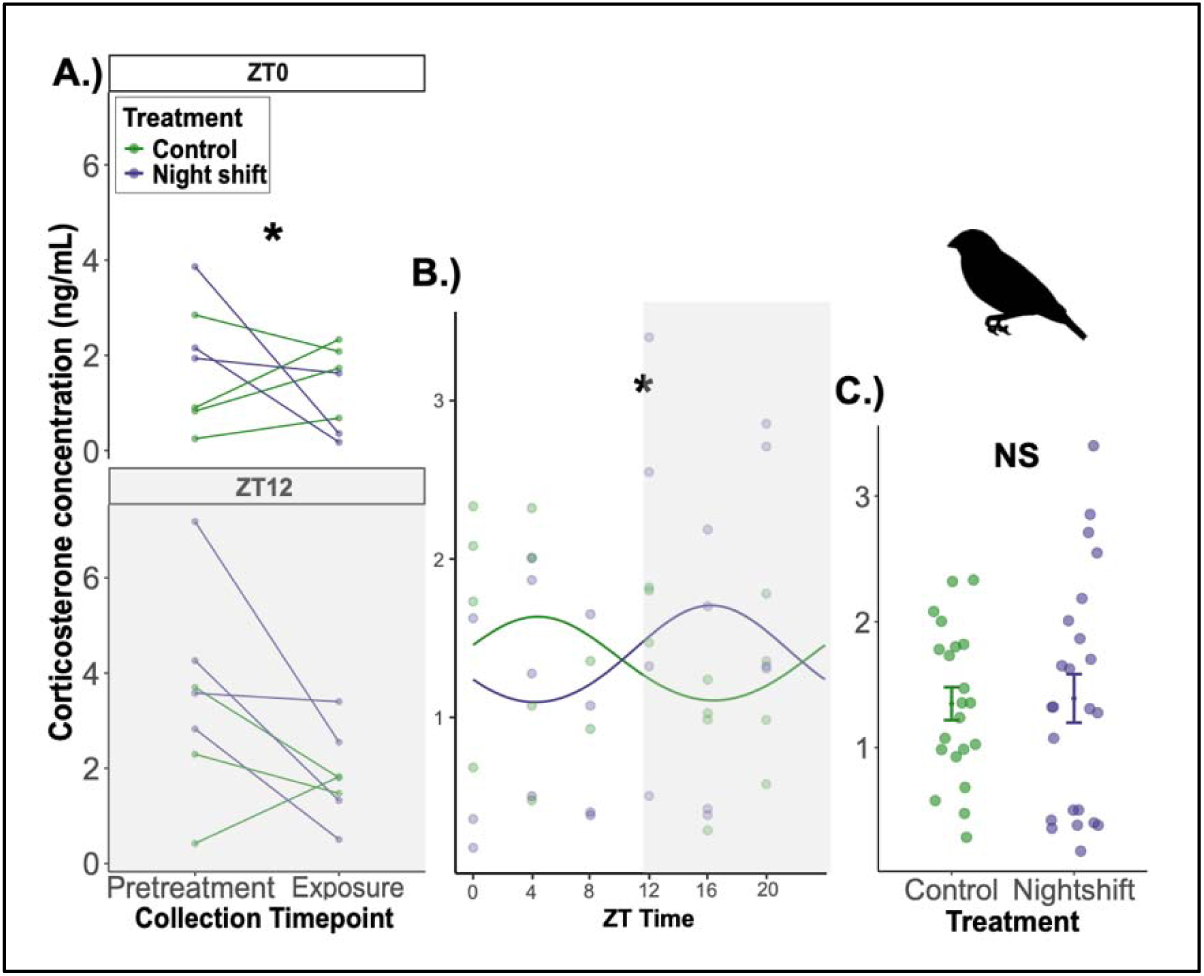
Corticosterone levels regarding circadian variation. A.) Corticosterone levels within birds before and after exposure simulated night shift work at specific ZT collection timepoints; B.) across 24-h period at exposure only, and C.) between treatment groups at exposure only. In all graphs, controls are depicted in green, while birds exposed to simulated night shift work are depicted in blue. A.) The X-axis indicates collection timepoint, while the Y-axis indicates corticosterone levels tracked between each unique individual before and after exposure to treatment. B.) The X-axis depicts the six ZT collection timeopints in which corticosterone levels were quantified. A sine wave is fit using GLMMcosine. C.) The X-axis indicates treatment groups, with average levels of corticosterone on the Y-axis using raw data on exposure data only Error bars indicate the standard error of the mean (SEM). Gray boxes indicate the dark phase. Asterisks denote statistical differences; \**P* < 0.05.

We found that both groups had significant corticosterone rhythms, indicated by the mesor (Control: GLMMcosinor: β = 0.30, 95% CI = 0.03 to 0.56; *p* = 0.03; Night shift: GLMMcosinor: β = 0.31, 95% CI = 0.07 to 0.56; *P* = 0.01; **Fig. 2B**). Moreover, both groups had relatively the same amplitude (Control: β = 0.19; Night shift: β = 0.22), with no statistical difference (GLMMcosinor: β = 0.03; *P* = 0.92) and no average difference in corticosterone levels (**Fig. 2C)**. We found that the acrophase of Cort peaked for control birds 4.49 hours from ZT0, resulting in the maximal level of Cort secreted approximately ∼ZT4-5 (10am-11am) (GLMMcosinor: β = 1.15 radians). In contrast, birds under night shift had their peak in Cort −7.76 hours from ZT24, resulting in the highest level of Cort approximately ∼ZT 16-17 (10pm-11pm) (GLMMcosinor: β = −2.03). Furthermore, there was statistically significant difference in the acrophase between groups due to simulated night shift treatment (GLMMcosinor: β = 3.09; 95% CI = 0.62 to 5.56; *P* = 0.01; **Figure 2B**).

Both groups had significant melatonin rhythms, indicated by the mesor (Control: GLMMcosinor: β = 157.43; 95% CI = 140.08 to 174.77; *P* < 0.0001; Night shift: GLMMcosinor: β = 171.57, 95% CI = 153.17 to 189.97; *P* < 0.0001; **Fig. 3A**). Both groups had similar amplitude estimates (Control: β = 17.31; Night shift: β = 10.72), with no statistically significant difference (GLMMcosinor: β = −6.58; *P* > 0.72) and no average difference in melatonin levels (**Fig. 3B)**. We found that the acrophase of melatonin release for control birds was −8.25 hours from ZT 24, resulting in the highest level of melatonin approximately ∼ZT 15-16 (9pm-10pm) (GLMMcosinor: β = −2.61). In contrast, the acrophase of melatonin for night shift birds was −0.65 hours from ZT 24, resulting in the highest level of melatonin observed approximately ∼ZT 23-24 (5am-6am) (GLMMcosinor: β = −0.18). We detected a slight trend indicating that the acrophase was different between groups, but this did not reach statistical significance (GLMMcosinor: β = 2.44; 95% CI = −0.38 to 5.26 ± 95% CI; *P* = 0.09)

**Figure 3.**
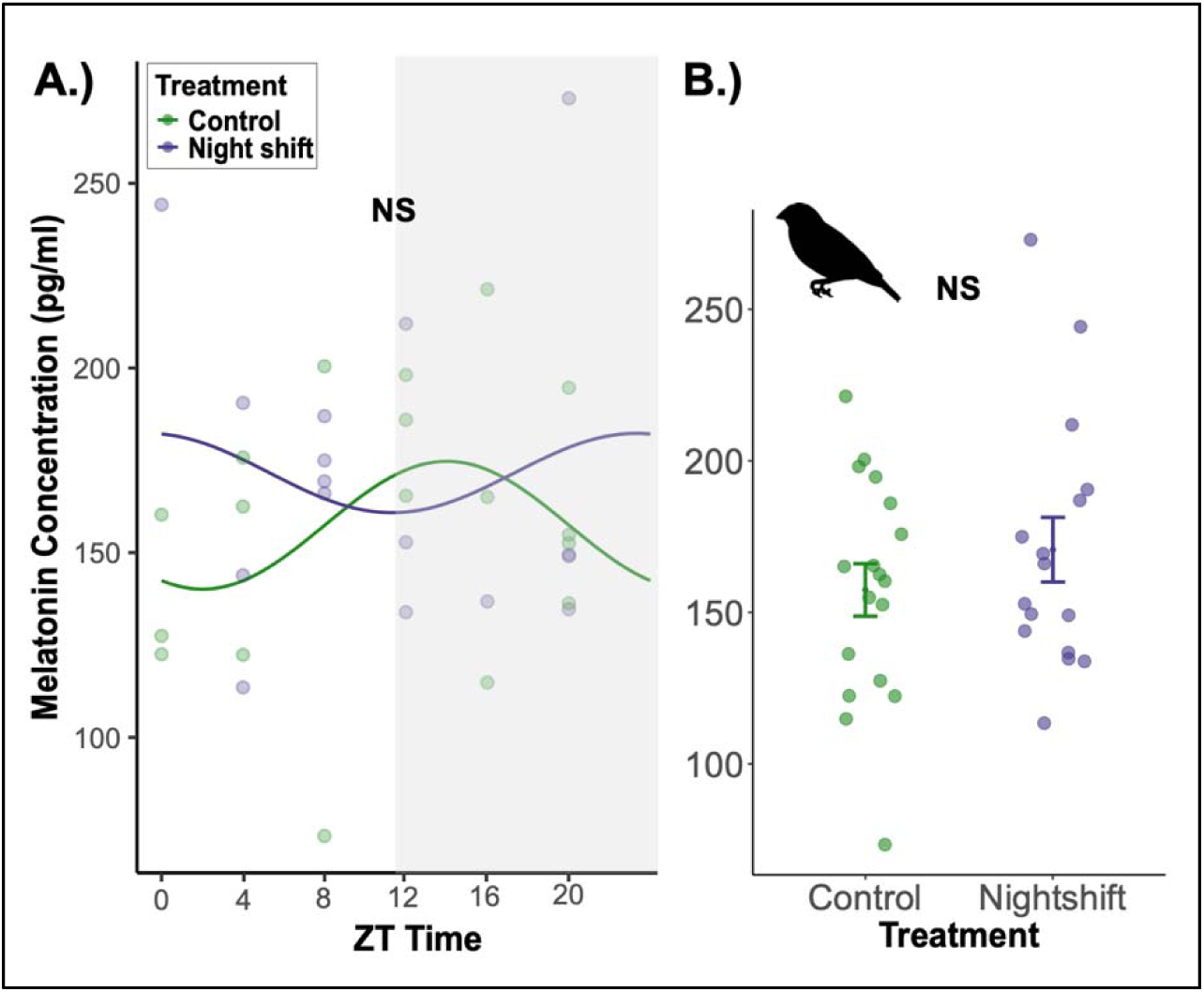
Melatonin rhythms and average hormone levels after exposure to night shift work. In all graphs, controls are depicted in green, while birds exposed to simulated night shift work are shown in blue. A.) The X-axis indicates six ZT collection times in which melatonin levels were quantified, while the Y-axis indicates melatonin levels (pg/mL). A best fit sine wave is fitted using GLMMcosine. B.) The X-axis depicts treatment group with the y-axis indicating melatonin levels. Average levels are plotted using raw data with error bars representing the standard error of the mean (SEM). Melatonin rhythms were significant, however, there was no difference in the acrophase or average difference in hormone levels between treatment groups.

### HPA axis reactivity

We did not detect a significant interaction between treatment and timepoint on post-restraint corticosterone levels (Treatment*Timepoint: F_2,75_ = 0.57; *P =* 0.57). However, there was an overall, significant main effect of treatment (F_1,38_ = 12.72; *P* < 0.001) in which post-restraint corticosterone levels in birds undergoing simulated night shift work were on average 2.53 ng/mL lower than those under a consistent light/dark cycle. (LMM: β = −2.53; 95% CI = −4.63 to −0.43; *t* = −2.29; *P =* 0.02; **Fig. 4**). There was no main effect of treatment (F_1,42_ = 0.84; *P =* 0.36) or its interaction with timepoint (F_2,63_ = 1.06; *P =* 0.35) on negative feedback efficiency. There was strong evidence indicating that there was an overall main effect of timepoint (F_2,63_ = 5.15; *P* ≤ 0.008) on negative feedback efficiency, however corticosterone levels did not differ when comparing between timepoints of interest (Pretreatment vs Acute: *P =* 0.21; Pretreatment vs Chronic: *P =* 0.34 **Fig. S2 and Fig. S3**)

**Figure 4.**
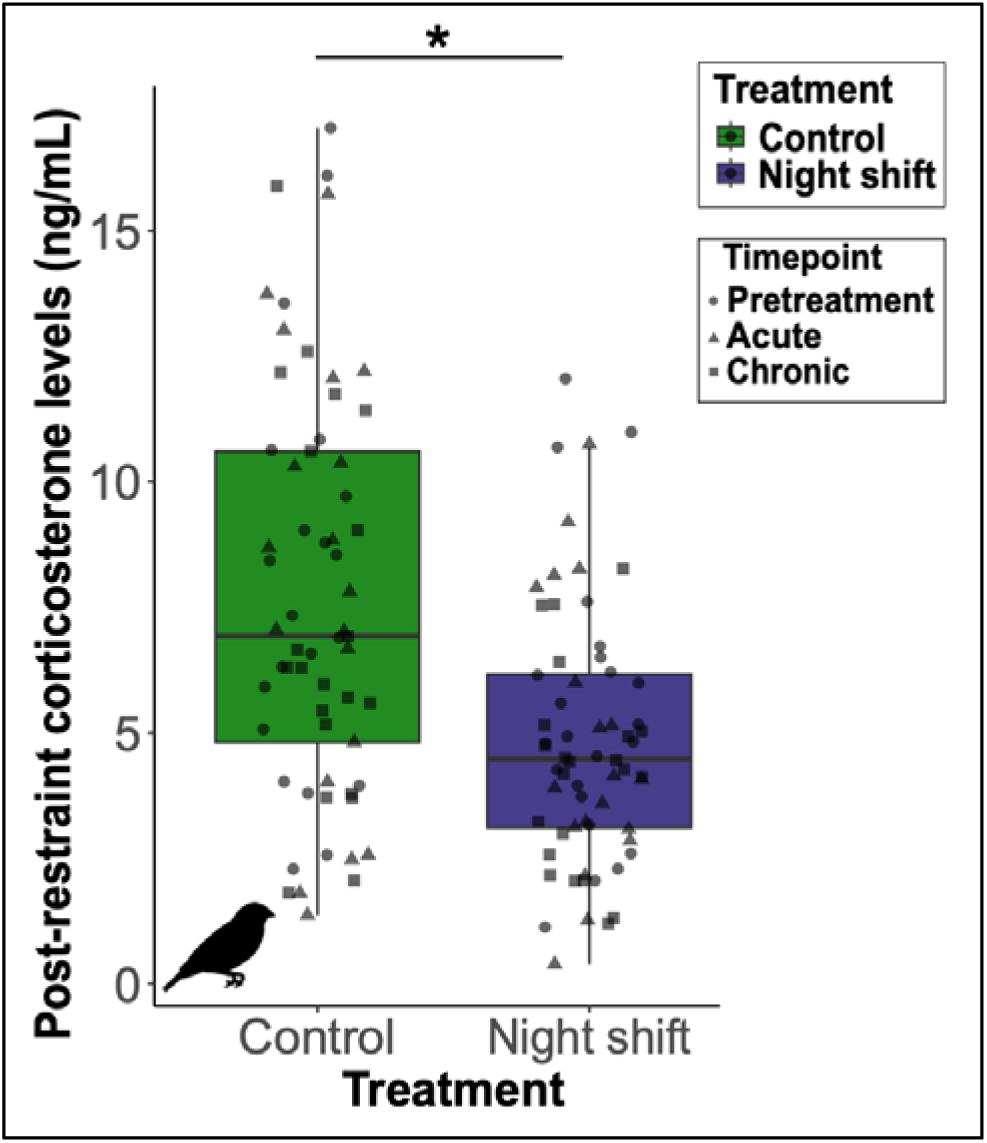
30 min post-restraint corticosterone levels between treatment groups represented as a box-and-whisker plot. The solid black line indicates the median value, with the upper 75% quartile and lower 25% quartile depicted, and the interquartile representing 50% of the observed data. The X-axis depicts our experimental groups, with control birds represented by the green bar and night shift birds represented by the blue bar. The Y-axis depicts corticosterone levels measured in ng/mL. Timepoints across the experiment are represented in shapes. Circle = Pretreatment, triangle = 3 night shifts, and square = 21 night shifts. Statistical significance is denoted with a solid black line with a single asterisk, with an associated *P-v*alue < 0.05).

### Morphometrics

We did not detect an overall main effect of treatment (F_1,41_ = 0.33; *P =* 0.57) or its interaction with timepoint (F_3,126_ = 1.65; *P =* 0.18) on body mass (**Fig S4**). There was no main effect of timepoint (X^2^ = 1.9; *P* = 0.59), nor the interaction between treatment or timepoint (X^2^ = 2.7; *P* = 0.44).

### Metabolites

We detected a significant interaction between Treatment and Timepoint, in which β-hydroxybutyrate levels differed between treatment groups but was dependent on the duration of treatment (F_2,83_ = 7.78; *P* = 0.0008). Post hoc tests revealed that there was no initial average difference between control and night shift groups in circulating β-hydroxybutyrate levels (Tukey’s Post hoc: β = 0.07; 95% CI = −0.70 to 0.83; *t* = 0.71; *P* = 0.86; **Figure 5A**). However, after acute exposure to night shift work (6 night shifts), treatment birds had on average 1.28 mmol/L greater amount of β-hydroxybutyrate than controls (Tukey’s Post hoc: β = 1.28; 95% CI = 0.53 to 2.04; *t* = 3.38; *P* = 0.0011; **Figure 5A**) and remained elevated until day 47 of the simulated night shift (Tukey’s Post hoc: β = 1.14; 95% CI = 0.39 to 1.9; *t* = 3.02; *P* = 0.004; **Figure 5A**).

**Figure 5.**
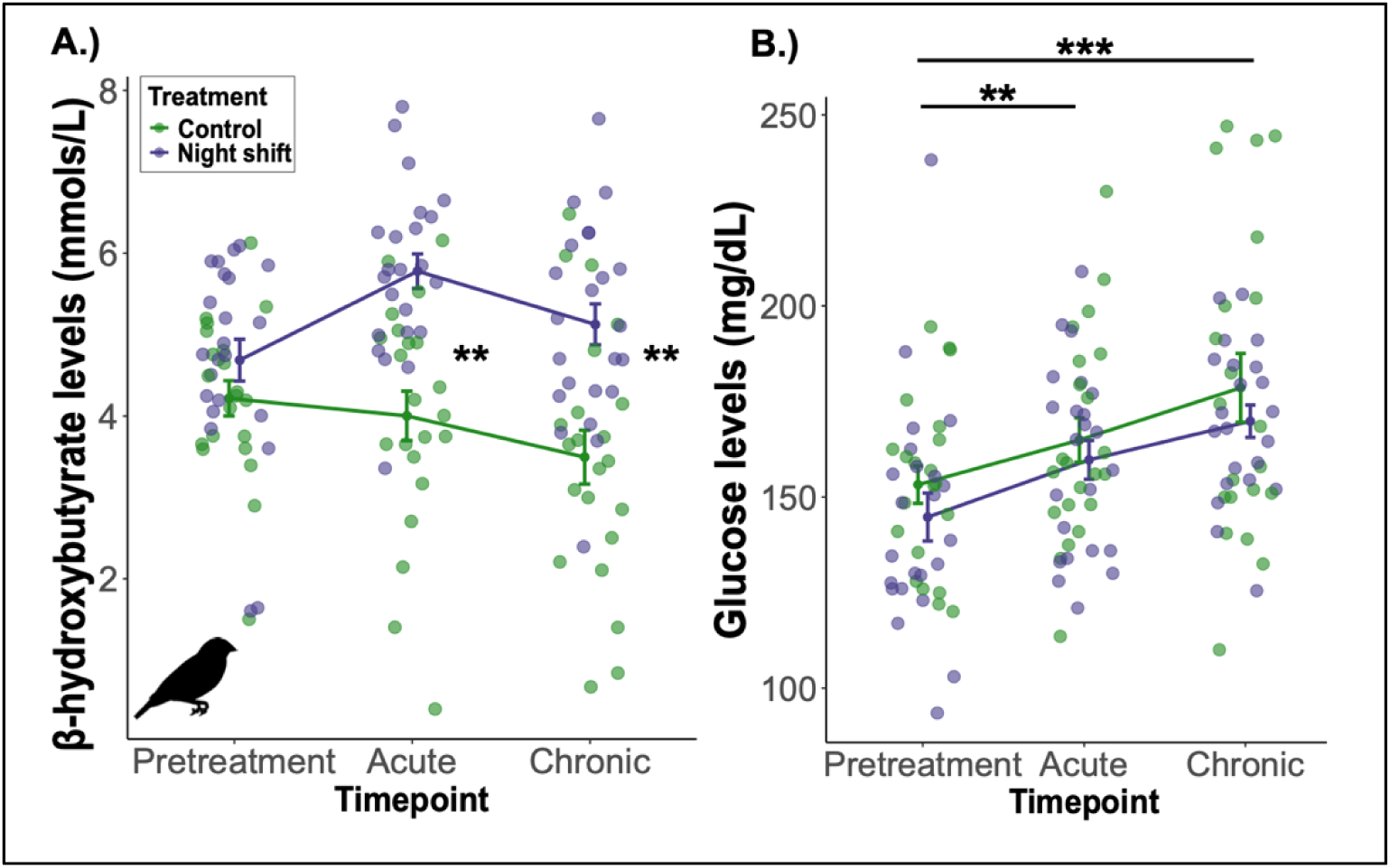
β-hydroxybutyrate and glucose levels between experimental groups over the duration of the experiment. For both graphs, the green line indicates controls, while the blue line indicates treatment birds. Averages are plotted using raw data, with error bars indicating the standard error of the mean (SEM). A.) The X-axis indicates Timepoint, while the Y-axis indicates circulating β-hydroxybutyrate levels (mmols/L). B.) The X-axis indicates timepoint, while the Y-axis indicates circulating glucose levels (mg/dL). Asterisks denote statistical differences; \*\**P* < 0.01, \*\*\**P* < 0.001.

We detected a significant main effect of Timepoint, in which glucose levels increased over the duration of the experiment (F_2,85_ = 11.8; *P* < 0.0001), however this was regardless of treatment, as there was no significant Treatment*Timepoint interaction (F_2,83_ = 0.10; *P* = 0.90). Specifically, glucose levels increased on average 13.24 mg/dL between Pretreatment and when birds were remeasured on day 11 of the experiment (acute timepoint) (LMM: β = 13.24; 95% CI = 3.10 to 23.32; *t* = 2.56; *P* < 0.01; **Fig. 3B**). Moreover, when comparing Pretreatment to day 47 of the experiment (chronic timepoint), we found that glucose increased on average 25.40 mg/dL more between the two timepoints (LMM: β = 25.40; 95% CI = 15.13 to 35.63; *t* = 4.85; *P* < 0.0001; **Fig. 3B**).

### Oxidative stress markers in the brain and liver

In the brain, there was a significant effect of treatment (F_1,29_ = 17.41; *P* = 0.0002) in which birds under night shift work had lower relative protein abundance of SOD1 compared to controls (LMM: β = −0.16; 95% CI = −0.24 to −0.08; *t* = −4.17; *P* = 0.0002; **Fig. 6A**). Although birds under night shift work had a reduction in SOD1 protein levels in the brain, this did not result in greater oxidative damage to lipids measured through 4-HNE adducts (F_1,30_ = 0.71; *p* = 0.41; **Fig 6B**). In the liver, there was no effect of treatment (F_1,37_ = 2.16; *p* = 0.15; **Fig. S5A**) on SOD1 levels or 4-HNE (F_1,36_ = 1.08; *p* = 0.31; **Fig. S5B**).

**Figure 6.**
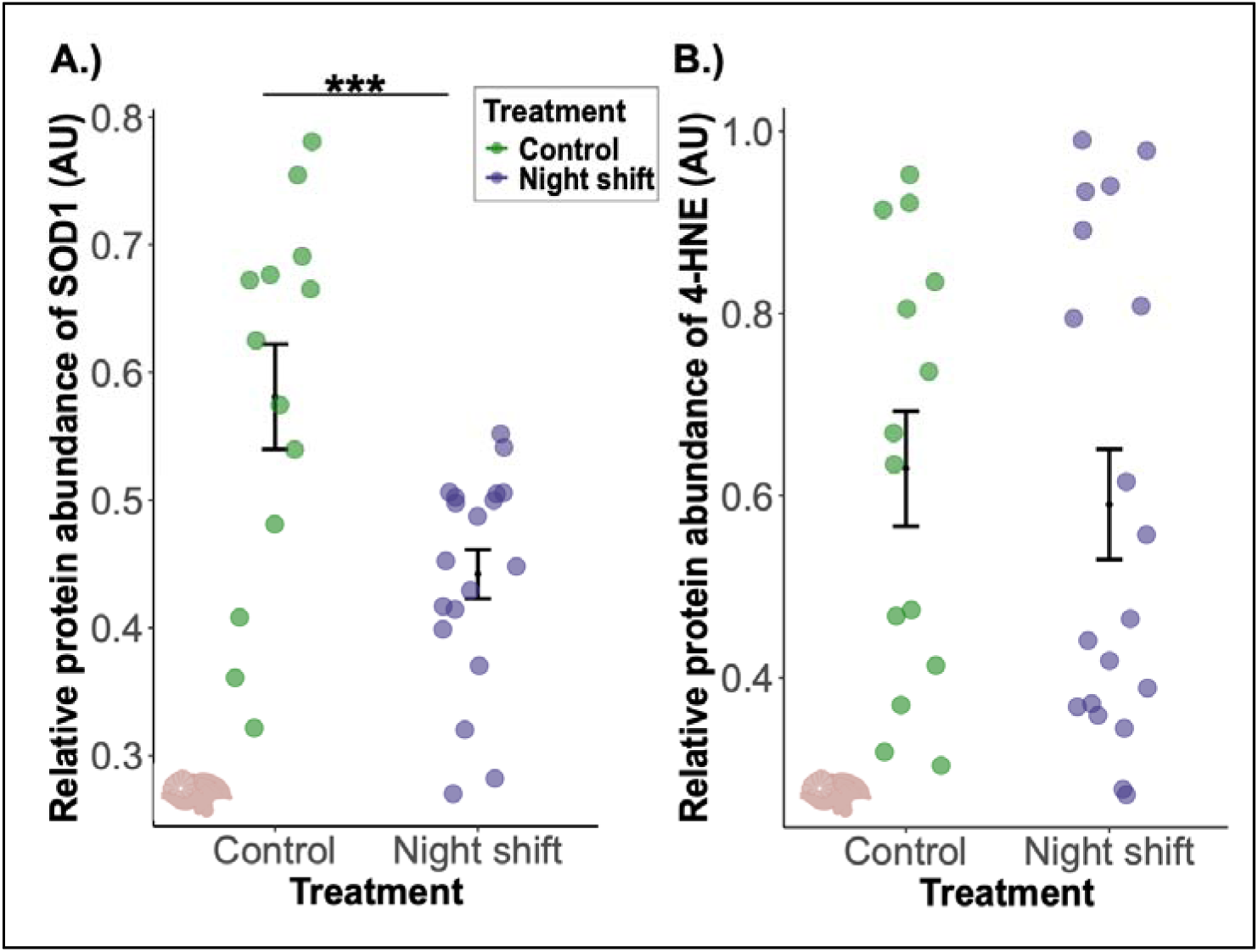
Relative abundance of SOD1 and 4-HNE protein levels in the brain. In both graphs, controls are depicted by the blue line, while birds exposed to night shift work are shown by the green line. The X-axis depicts treatment groups, while the Y-axis indicates relative protein abundance. A.) Sample size for this analysis was n = 13 for controls, and n = 19 for treatment birds. B.) Sample size for this analysis was n = 14 for controls, and n = 19 for treatment birds. Averages are plotted using raw data, with error bars indicating the standard error of the mean (SEM). Note that we show untransformed data, while in the statistical analysis, we log transform both response variables. Asterisks with lines denote statistical differences; *\*P <* 0.05*, **P <* 0.01*, ***P* < 0.001.

## Discussion

In this study, we examined how simulated night shift work altered corticosterone and melatonin rhythms resulting in changes in HPA axis dynamics and downstream consequences on energy metabolism, body condition, and oxidative balance. We found that simulated night shift inverted corticosterone profiles, with peak levels detected during the dark phase between 10-11pm, rather than during the light phase. Similarly, there was a slight trend indicating a shift in the acrophase of melatonin towards the early morning hours, rather than at night. Plasma constraints limited our ability to measure melatonin from samples collected towards the beginning of the light phase (ZT0) and the middle of the night (ZT16), likely contributing to inadequate variation to detect a stronger (or weaker) shift in the acrophase. Nevertheless, we still found significant melatonin and corticosterone rhythms in birds undergoing simulated night shift work, suggesting that they were able to tolerate the treatment paradigm and maintain circadian alignment, albeit inverted from their natural diurnal rhythm.

The inversion of corticosterone profiles may be a plastic response to alternating patterns of light at night, in which circadian oscillators resynchronize to predictable nighttime conditions to prevent a mismatch between endogenous rhythms - yet may not be without some cost. In our study, the inversion of corticosterone profiles altered physiological responses and energy homeostasis. Birds under simulated night shift work had a dampened glucocorticoid response to capture and restraint, indicating poor HPA axis reactivity. Furthermore, there was a significant reduction in corticosterone profiles at ZT0 (lights on; “awakening”) after exposure to treatment.

Poor sleep quality, a common outcome due to consistent/rotating night shift work in humans, show conflicting patterns on stress responsiveness, with studies indicating a suppressed or heightened response (van Dalfsen and Markus, 2018). This variation may stem from the fact that different aspects of sleep (*i.e.,* duration, disturbance, or architecture) can differentially affect neuroendocrine responses, along with other factors such as sex (Bassett et al., 2015). However, the strongest predictor that seems to be associated with the blunting of stress responsiveness in humans is daytime sleepiness (van Dalfsen and Markus, 2018). In our study, it is possible that birds under night shift work were unable to respond strongly to capture and restraint due to the fact that they were previously under the treatment paradigm the morning that we tested reactivity of the HPA axis (*i.e.,* altered activity levels or loss of sleep). Given that corticosterone profiles were inverted and strongly rhythmic, it is possible behavioral rhythms followed in parallel, as there is tight coupling between both types of rhythms (Oster et al., 2017; Rao and Androulakis, 2019). However, we are unable to confirm this speculation as we did not measure behavioral rhythms or sleep patterns and it is possible for glucocorticoid and behavioral rhythms to disassociate through masking effects.

Although stress responsiveness was not tested, a similar inversion of glucocorticoids profiles (higher levels at night, rather than the morning) measured in saliva was found in female nurses working consistent night shifts (Niu et al., 2015). Moreover, several studies corroborate that salivary cortisol levels at the time of awakening are significantly lower in individuals working night shifts (Andreadi et al., 2025; Hennig et al., 1998; Niu et al., 2015). Interestingly, Niu et al., 2015 found that salivary cortisol levels remained lower in night shift nurses, even on the first day off the shift schedule, indicating a lingering effect. However, cortisol levels rebounded and were not significantly different from day shift nurses by the 2^nd^ day off night shift. This finding further suggests plasticity of the circadian system to quickly realign to a newly imposed pattern of wakefulness/rest in a relatively short amount of time. In our study, we measured 24-h corticosterone profiles on the first day after three simulated night shifts to capture the strongest effect of our treatment paradigm. Our results may have produced similar results as Niu et al., 2015 if we had sampled 24-h hormone profiles on the second day off simulated night shift after allowing birds the opportunity to readjust to a different pattern (12L:12D). This reorganization between circadian oscillators suggests high flexibility to disrupting patterns of light at night. Yet, there are likely species-specific differences in the time it takes to re-entrain to a light/dark cycle based on sensitivity to photoperiodism or the presence of supplementary environmental cues.

For birds (and mammals) within temperate zones that heavily rely on photoperiodism as an ultimate cue, circadian oscillators may be slower or more inflexible in their response to phase changes (*i.*e., days getting longer/shorter) given that they need to subsequently finetune downstream responses with proximal cues (*e.g.,* food availability/temperature) to prevent mismatches (Tolla and Stevenson, 2020). One study found significantly more variation in the photoperiodic response of opportunistic breeders compared to seasonal breeders (measured through gonadal stimulation/growth), indicating that seasonal breeders, to some extent, lack flexibility in response to photoperiodism and it is more of a fixed response (Watts et al., 2015). A study in Siberian hamster (seasonal breeder) found that supplementary cues (food availability and population size) became more important to time reproductive events when they were placed in a photoperiod unable to activate reproductive pathways, indicating flexibility when photoperiod is no longer sufficient or predictable (Paul et al., 2009). However, circadian genes or hormone profiles were not measured in either study. Nevertheless, our finding on inverted corticosterone profiles in response to simulated night shift work suggests that a diurnal, opportunistic songbird, can quickly adapt to unnatural patterns of light at night and could affect how they coordinate peripheral or master oscillators to maintain function.

Glucose metabolism is tightly regulated such that animals do not exceed or fall short of optimal ranges that risks hyper/hypoglycemia. Elevated blood glucose levels glycate biomolecules, increase oxidative stress markers, and damage epithelial lining of blood vessels in most animals (Preiser et al., 2016). Consequently, hyperglycemia is strongly associated with the development of atherosclerosis, hypertension, and insulin insensitivity, all of which are risk factors underlying the increased prevalence of cardiovascular diseases in night shift workers (Briançon-Marjollet et al., 2015; James et al., 2017). Rodent models of shift work and other perturbations in the nighttime light environment also show similar metabolic consequences as humans, such as elevated fasted blood glucose levels, and a reduction in insulin sensitivity (Opperhuizen et al., 2015). In our study, we did not detect changes in fasted blood glucose levels due to night shift work. This is likely due to the fact that avian species are able to withstand a wide range of glucose values, which reflect variation in environmental pressures, nutritional state, life-history, or seasons (Beattie et al., 2022; Remage-Healey and Romero, 2000). However, we did see an interesting overall increase in fasted glucose values throughout the duration of the experiment, which may be an indicator of repeated sampling and handling stress. Montoya et al., (2020) found that glucose levels increased due to successive handling to administer intraperitoneal injections within a much shorter timeframe (0-40 minutes) in zebra finches (Montoya et al., 2020). In a previous experiment using a constant light paradigm in zebra finches, we found a similar increase in baseline glucose levels in control birds’ overtime, however, it was not statistically significant (Pham et al., 2025). In this study, we sampled over a much longer duration (days), yet still found an increase of glucose levels over time in both groups.

This could be a plastic response to modify the setpoint of circulating glucose levels to appropriately respond to predictable stressors, such as repeated handling/injections. While baseline glucose levels were unaltered in response to inclement weather, adult tree swallows that previously experienced colder temperatures mounted a stronger glucose response to restraint stress compared to swallows that experienced warmer temperatures (Ryan et al., 2023). Thus, changes in glucose regulation could prime individuals to adaptively respond to predictable challenges they may encounter in the future.

Birds under night shift lighting patterns had significantly higher β-hydroxybutyrate metabolites compared to controls. This finding suggests that they were under a greater energetic demand to metabolize fats for fuel, rather than for storage. Given that we found inverted corticosterone rhythms, this is somewhat paradoxical as anabolism should be favored in parallel to corticosterone rhythms. β-hydroxybutyrate levels are generally only detected when glucose is no longer the primary substrate for energy production (*i.e.,* fasting conditions) (Tabatabaei Dakhili et al., 2025). Within the context of our study, we fasted our birds to eliminate sources of variation due to food consumption prior to sampling, but also to evaluate if night shift work differentially affected metabolic pathways. Indeed, the higher β-hydroxybutyrate levels detected in night shift birds could be suggestive of a compensatory mechanism to maintain sufficient energy and activate cellular pathways in costly tissues, such as the brain or cardiac muscle (Tabatabaei Dakhili et al., 2025). Ketone bodies activate antioxidant pathways (among others), suggesting that under greater energetic demands or fasting, higher breakdown of lipids (*i.e.,* higher ketone bodies) may preserve or enhance brain and cardiac function through activating cellular stress response pathways (Rojas-Morales et al., 2020; Tabatabaei Dakhili et al., 2025).

This perspective and our finding of higher β-hydroxybutyrate levels could explain the patterns detected regarding our oxidative stress markers. Simulated night shift work decreased SOD1 protein levels in the brain, but birds did not display elevated oxidative damage to lipids. Furthermore, there were no changes in SOD or 4-HNE in the liver, indicating a tissue-specific response of oxidative stress under night shift work. The decrease in SOD1 levels in the brain could suggest a utilization, in which SOD1 was depleted in order to convert superoxide into hydrogen peroxide. Because β-hydroxybutyrate can activate antioxidant pathways, we would expect to see greater levels of SOD1 in the brain, but this was not the case. Given the opposing patterns that we detected with regards to oxidative damage in the brain, perhaps there was a counter-balancing effect, in which SOD1 levels were continuously depleted and replenished via β-hydroxybutyrate signaling to prevent the accumulation of damage.

Contrary to our predictions, birds under night shift work did not have a suppression in their melatonin levels. This finding may further explain why we did not detect elevated oxidative damage because melatonin can quench reactive species. However, we only measured one marker of antioxidants and oxidative damage. It is possible night shift work differentially affects biomolecules, with some more vulnerable than others or repaired at different rates. In other studies, melatonin supplementation can realign circadian oscillators after jet lag conditions and reduce accumulation of oxidative damage, likely by quenching a wide range of reactive species (Bonnefont-Rousselot and Collin, 2010; Zisapel, 2018).

While circadian misalignment is detrimental in many cases, through masking, diurnal and nocturnal animals may be able to unlock temporal niches that do not override circadian-gated mechanisms. For example, although zebra finches exposed to ecologically relevant levels of artificial light at night displayed higher activity bouts at night (indicating some loss of sleep), circadian gene regulation was maintained in the brain (Alaasam et al., 2021). Moreover, after 6 months of light at night, zebra finches were able to habituate and did not exhibit severe levels of oxidative stress (Alaasam et al., 2024). Therefore, this is an example of positive behavioral masking, with seemingly little physiological costs associated with the masking response.

In a mammalian model, *Octodon degus* are able to leverage both temporal niches. While degus are mainly diurnal, they can display inverted phases in activity and body temperatures compared to diurnal chronotypes (Otalora et al., 2010). Yet, other parameters such as neuropeptides associated with awakening, and core clock genes are invariable between diurnal and nocturnal degus (Otalora et al., 2010). Therefore, the ability to leverage the changes in the nighttime environment through behavioral or physiological masking, without overriding circadian-gated mechanisms may be an important plastic response in the face of modified ecological landscapes and modern lifestyles (Bahammam, 2025; Helm et al., 2017).

## Conclusions

Corticosterone and melatonin are reliable indicators of circadian alignment. Thus, it is striking to see that birds maintained significant rhythms after 66 days of simulated night shift lighting patterns. Such minimal effects of our treatment paradigm on body condition and oxidative damage suggests that birds may have been able to predict and habituate to the switches in lighting patterns by quickly realigning themselves to the imposed change through plastic mechanisms. These explanations seem plausible, especially because of the aforementioned study in nurses demonstrating that by the second day off night shift, cortisol patterns returned to levels similar to those working day shifts (Niu et al., 2015). This is the first study to our knowledge to describe similar corticosterone profile changes in human night shift workers using a diurnal songbird as the model system. Moreover, we extend upon previous findings, by showing that zebra finches display even more circadian plasticity through sustained, albeit inverted, hormone profiles in response to chronic, simulated night shift work.

## Acknowledgments

The authors would like to thank the Wada/Hood/Hill labs for helpful comments on improving the manuscript. The authors would also like to thank the undergraduate researchers, Yeojin Joo, Darielys Diez, Jordan Shapach, and Katie Gross as well as graduate students in the Wada lab for their assistance with animal husbandry, experimental setup, and sample collection. Lastly, the authors would like to thank Dr.’s Jenny Ouyang and Valentina Alaasam for helpful comments, suggestions, and sharing their melatonin assay protocol.

## Citations

Alaasam, V.J., Duncan, R., Casagrande, S., Davies, S., Sidher, A., Seymoure, B., Shen, Y., Zhang, Y., Ouyang, J.Q., 2018. Light at night disrupts nocturnal rest and elevates glucocorticoids at cool color temperatures. J. Exp. Zool. Part A Ecol. Integr. Physiol. 329, 465–472. 10.1002/jez.2168

Alaasam, V.J., Hui, C., Lomas, J., Ferguson, S.M., Zhang, Y., Yim, W.C., Ouyang, J.Q., 2024. What happens when the lights are left on? Transcriptomic and phenotypic habituation to light pollution. iScience 27, 108864. 10.1016/j.isci.2024.108864

Alaasam, V.J., Liu, X., Niu, Y., Habibian, J.S., Pieraut, S., Ferguson, B.S., Zhang, Y., Ouyang, J.Q., 2021. Effects of dim artificial light at night on locomotor activity, cardiovascular physiology, and circadian clock genes in a diurnal songbird. Environ. Pollut. 282, 117036. 10.1016/j.envpol.2021.117036

Andreadi, A., Andreadi, S., Todaro, F., Ippoliti, L., Bellia, A., Magrini, A., Chrousos, G.P., Lauro, D., 2025. Modified Cortisol Circadian Rhythm: The Hidden Toll of Night-Shift Work. Int. J. Mol. Sci. 26, 1–16. 10.3390/ijms26052090

Asiamah, N., Mends-Brew, E., Boison, B.K.T., 2021. A spotlight on cross-sectional research: Addressing the issues of confounding and adjustment. Int. J. Healthc. Manag. 14, 183–196. 10.1080/20479700.2019.1621022

Bahammam, A.S., 2025. From Wings to Wellness: A Research Agenda Inspired by Migratory Bird Adaptations for Sleep and Circadian Medicine. Nat. Sci. Sleep 17, 583–595. 10.2147/NSS.S519493

Bassett, S.M., Lupis, S.B., Gianferante, D., Rohleder, N., Wolf, J.M., 2015. Sleep quality but not sleep quantity effects on cortisol responses to acute psychosocial stress. Stress 18, 638–644. 10.3109/10253890.2015.1087503

Beattie, U.K., Ysrael, M.C., Lok, S.E., Romero, L.M., 2022. The Effect of a Combined Fast and Chronic Stress on Body Mass, Blood Metabolites, Corticosterone, and Behavior in House Sparrows (Passer domesticus). Yale J. Biol. Med. 95, 19–31.

Bedrosian, T.A., Fonken, L.K., Nelson, R.J., 2016. Endocrine Effects of Circadian Disruption. Annu. Rev. Physiol. 78, 109–131. 10.1146/annurev-physiol-021115-105102

Bonnefont-Rousselot, D., Collin, F., 2010. Melatonin: Action as antioxidant and potential applications in human disease and aging. Toxicology 278, 55–67. 10.1016/j.tox.2010.04.008

Braun, E.J., Sweazea, K.L., 2008. Glucose regulation in birds. Comp. Biochem. Physiol. -B Biochem. Mol. Biol. 151, 1–9. 10.1016/j.cbpb.2008.05.007

Briançon-Marjollet, A., Weiszenstein, M., Henri, M., Thomas, A., Godin-Ribuot, D., Polak, J., 2015. The impact of sleep disorders on glucose metabolism: Endocrine and molecular mechanisms. Diabetol. Metab. Syndr. 7, 1–16. 10.1186/s13098-015-0018-3

Brun, C., Hernandez-Alba, O., Hovasse, A., Criscuolo, F., Schaeffer-Reiss, C., Bertile, F., 2022. Resistance to glycation in the zebra finch: Mass spectrometry-based analysis and its perspectives for evolutionary studies of aging. Exp. Gerontol. 164. 10.1016/j.exger.2022.111811

Caruso, C.C., 2014. Negative impacts of shiftwork and long work hours. Rehabil. Nurs. 39, 16–25. 10.1002/rnj.107

Cassone, V.M., 2015. Avian circadian organization. Mech. Circadian Syst. Anim. Their Clin. Relev. 35, 69–94. 10.1007/978-3-319-08945-4_5

Cassone, V.M., 2014. Avian circadian organization: A chorus of clocks. Front. Neuroendocrinol. 35, 76–88. 10.1016/j.yfrne.2013.10.002

Costa, G., 2003. Shift work and occupational medicine: An overview. Occup. Med. (Chic. Ill). 53, 83–88. 10.1093/occmed/kqg045

Cox, D.T.C., Gaston, K.J., 2024. Cathemerality: a key temporal niche. Biol. Rev. 99, 329–347. 10.1111/brv.13024

Guindon, G.E., Murphy, C.A., Milano, M.E., Seggio, J.A., 2024. Turn off that night light! Light-at-night as a stressor for adolescents. Front. Neurosci. 18, 1–8. 10.3389/fnins.2024.1451219

Hansen, J., Pedersen, J.E., 2025. Night shift work and breast cancer risk – 2023 update of epidemiologic evidence. J. Natl. Cancer Cent. 5, 94–103. 10.1016/j.jncc.2024.07.004

Helm, B., Visser, M.E., Schwartz, W., Kronfeld-Schor, N., Gerkema, M., Piersma, T., Bloch, G., 2017. Two sides of a coin: Ecological and chronobiological perspectives of timing in the wild. Philos. Trans. R. Soc. B Biol. Sci. 372. 10.1098/rstb.2016.0246

Hennig, J., Kieferdorf, P., Moritz, C., Huwe, S., Netter, P., 1998. Changes in cortisol secretion during shiftwork: Implications for tolerance to shiftwork? Ergonomics 41, 610–621. 10.1080/001401398186784

Husse, J., Kiehn, J.T., Barclay, J.L., Naujokat, N., Meyer-Kovac, J., Lehnert, H., Oster, H., 2017. Tissue-specific dissociation of diurnal transcriptome rhythms during sleep restriction in mice. Sleep 40. 10.1093/sleep/zsx068

James, S.M., Honn, K.A., Gaddameedhi, S., Van Dongen, H.P.A., 2017. Shift Work: Disrupted Circadian Rhythms and Sleep—Implications for Health and Well-being. Curr. Sleep Med. Reports 3, 104–112. 10.1007/s40675-017-0071-6

Jimeno, B., Briga, M., Hau, M., Verhulst, S., 2018. Male but not female zebra finches with high plasma corticosterone have lower survival. Funct. Ecol. 32, 713–721. 10.1111/1365-2435.13021

Kervezee, L., Cuesta, M., Cermakian, N., Boivin, D.B., 2019. The Phase-Shifting Effect of Bright Light Exposure on Circadian Rhythmicity in the Human Transcriptome. J. Biol. Rhythms 34, 84–97. 10.1177/0748730418821776

Kriengwatana, B., Wada, H., Schmidt, K.L., Taves, M.D., Soma, K.K., MacDougall-Shackleton, S.A., 2014. Effects of nutritional stress during different developmental periods on song and the hypothalamic-pituitary-adrenal axis in zebra finches. Horm. Behav. 65, 285–293. 10.1016/j.yhbeh.2013.12.013

Mendoza, J., 2021. Nighttime Light Hurts Mammalian Physiology: What Diurnal Rodent Models Are Telling Us. Clocks and Sleep 3, 236–250. 10.3390/clockssleep3020014

Mishra, I., Knerr, R.M., Stewart, A.A., Payette, W.I., Richter, M.M., Ashley, N.T., 2019. Light at night disrupts diel patterns of cytokine gene expression and endocrine profiles in zebra finch (Taeniopygia guttata). Sci. Rep. 9, 1–12. 10.1038/s41598-019-51791-9

Monographs, I., Identification, O.N.T.H.E., Hazards, O.F.C., Humans, T.O., 2019. Carcinogenicity of night shift work, The Lancet. Oncology. 10.1016/S1470-2045(19)30455-3

Montoya, B., Briga, M., Jimeno, B., Verhulst, S., 2020. Glucose regulation is a repeatable trait affected by successive handling in zebra finches. J. Comp. Physiol. B Biochem. Syst. Environ. Physiol. 190, 455–464. 10.1007/s00360-020-01283-4

Moon, J., Ikeda-Araki, A., Mun, Y., 2024. Night shift work and female breast cancer: a two-stage dose-response meta-analysis for the correct risk definition. BMC Public Health 24, 1–16. 10.1186/s12889-024-19518-2

Niu, S.F., Chung, M.H., Chu, H., Tsai, J.C., Lin, C.C., Liao, Y.M., Ou, K.L., O’Brien, A.P., Chou, K.R., 2015. Differences in cortisol profiles and circadian adjustment time between nurses working night shifts and regular day shifts: A prospective longitudinal study. Int. J. Nurs. Stud. 52, 1193–1201. 10.1016/j.ijnurstu.2015.04.001

Opperhuizen, A.L., van Kerkhof, L.W.M., Proper, K.I., Rodenburg, W., Kalsbeek, A., 2015. Rodent models to study the metabolic effects of shiftwork in humans. Front. Pharmacol. 6, 1–20. 10.3389/fphar.2015.00050

Oster, H., Challet, E., Ott, V., Arvat, E., de Kloet, E.R., Dijk, D.J., Lightman, S., Vgontzas, A., Van Cauter, E., 2017. The functional and clinical significance of the 24-hour rhythm of circulating glucocorticoids. Endocr. Rev. 38, 3–45. 10.1210/er.2015-1080

Otalora, B.B., Vivanco, P., Madariaga, A.M., Madrid, J.A., Rol, M.Á., 2010. Internal temporal order in the circadian system of a dual-phasing rodent, the octodon degus. Chronobiol. Int. 27, 1564–1579. 10.3109/07420528.2010.503294

Ouyang, J.Q., Davies, S., Dominoni, D., 2018. Hormonally mediated effects of artificial light at night on behavior and fitness: Linking endocrine mechanisms with function. J. Exp. Biol. 221. 10.1242/jeb.156893

Paul, M.J., Galang, J., Schwartz, W.J., Prendergast, B.J., 2009. Intermediate-duration day lengths unmask reproductive responses to nonphotic environmental cues. Am. J. Physiol. - Regul. Integr. Comp. Physiol. 296, 1613–1619. 10.1152/ajpregu.91047.2008

Perfito, N., 2010. The reproductive and stress physiology of Zebra Finches in context: Integrating field and laboratory studies. Emu 110, 199–208. 10.1071/MU09091

Perfito, N., Kwong, J.M.Y., Bentley, G.E., Hau, M., 2008. Cue hierarchies and testicular development: Is food a more potent stimulus than day length in an opportunistic breeder (Taeniopygia g. guttata)? Horm. Behav. 53, 567–572. 10.1016/j.yhbeh.2008.01.002

Pilcher, J.J., Lambert, B.J., Huffcutt, A.I., 2000. Differential effects of permanent and rotating shifts on self-report sleep length: A meta-analytic review. Sleep 23, 155–163. 10.1093/sleep/23.2.1b

Preiser, J.C., Thooft, A., Tironi, R.M., 2016. Stress hyperglycemia. Stress Response Crit. Illn. Metab. Horm. Asp. 89–94. 10.1007/978-3-319-27687-8_8

Rao, R., Androulakis, I.P., 2019. The physiological significance of the circadian dynamics of the HPA axis: Interplay between circadian rhythms, allostasis and stress resilience. Horm. Behav. 110, 77–89. 10.1016/j.yhbeh.2019.02.018

Remage-Healey, L., Romero, L.M., 2000. Daily and Seasonal Variation in Response to Stress in Captive Starlings (Sturnus Vulgaris): Glucose. Gen. Comp. Endocrinol. 119, 60–68. 10.1006/gcen.2000.7492

Rojas-Morales, P., Pedraza-Chaverri, J., Tapia, E., 2020. Ketone bodies, stress response, and redox homeostasis. Redox Biol. 29, 101395. 10.1016/j.redox.2019.101395

Spierings, M.J., Cate, C. Ten, 2016. Zebra finches as a model species to understand the roots of rhythm. Front. Neurosci. 10, 1–3. 10.3389/fnins.2016.00345

Sweazea, K.L., 2022a. Revisiting glucose regulation in birds – A negative model of diabetes complications. Comp. Biochem. Physiol. Part - B Biochem. Mol. Biol. 262, 110778. 10.1016/j.cbpb.2022.110778

Sweazea, K.L., 2022b. Revisiting glucose regulation in birds – A negative model of diabetes complications. Comp. Biochem. Physiol. Part - B Biochem. Mol. Biol. 262, 23332973. 10.1016/j.cbpb.2022.110778

Tabatabaei Dakhili, S.A., Yang, K., Stenlund, M.J., Ussher, J.R., 2025. The multifaceted roles of ketones in physiology. Exp. Physiol. 1–13. 10.1113/EP092243

Tolla, E., Stevenson, T.J., 2020. Sex differences and the neuroendocrine regulation of seasonal reproduction by supplementary environmental cues. Integr. Comp. Biol. 60, 1506–1516. 10.1093/icb/icaa096

van Dalfsen, J.H., Markus, C.R., 2018. The influence of sleep on human hypothalamic– pituitary–adrenal (HPA) axis reactivity: A systematic review. Sleep Med. Rev. 39, 187–194. 10.1016/j.smrv.2017.10.002

Van, N.T.H., Hoang, T., Myung, S.K., 2021. Night shift work and breast cancer risk: A meta-analysis of observational epidemiological studies. Carcinogenesis 42, 1260– 1269. 10.1093/carcin/bgab074

Walker, W.H., Walton, J.C., DeVries, A.C., Nelson, R.J., 2020. Circadian rhythm disruption and mental health. Transl. Psychiatry 10. 10.1038/s41398-020-0694-0

Watts, H.E., Macdougall-Shackleton, S.A., Hahn, T.P., 2015. Variation among individuals in photoperiod responses: Effects of breeding schedule, photoperiod, and age-related photoperiodic experience in birds. J. Exp. Zool. Part A Ecol. Genet. Physiol. 323, 368–374. 10.1002/jez.1929

Zisapel, N., 2018. New perspectives on the role of melatonin in human sleep, circadian rhythms and their regulation. Br. J. Pharmacol. 175, 3190–3199. 10.1111/bph.14116

