## Supplementary Files for "Mimicking the night shift: Hypothalamic-pituitary-adrenal (HPA) axis regulation, energy homeostasis, and oxidative stress in a model diurnal songbird"

Running title: Simulated night shift works reduces stress reactivity and elevates lipid catabolism in zebra finches

Kevin Pham*^1^, Victoria M. Coutts^2^, Alexander J. Hoffman^1^, and Haruka Wada^1^

^1^Department of Biological Sciences, Auburn University, Auburn, AL 36849, USA

^2^Department of Biological Sciences, Louisiana State University, Baton Rouge, LO 36849, USA

**Precision Xtra device validation**

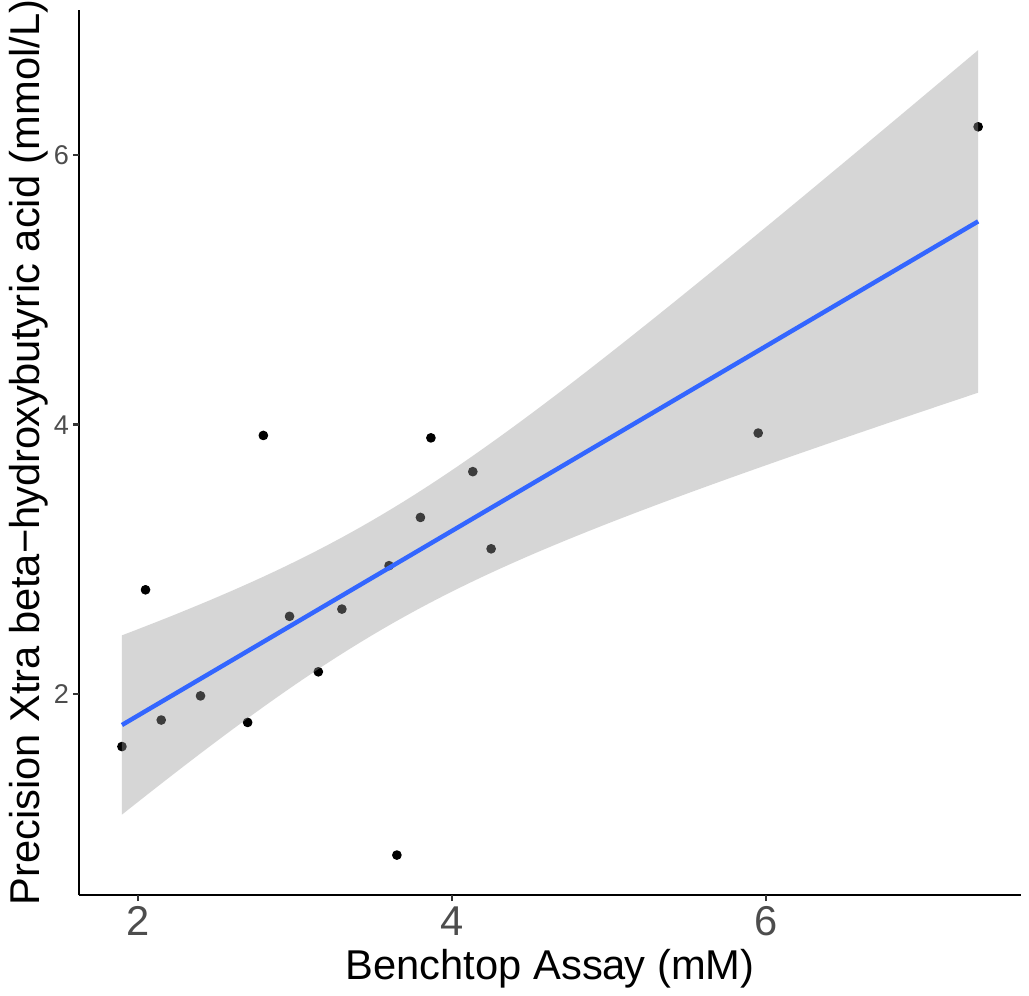
Male and female zebra finches (N = 13), which were not part of this study, were briefly fasted the night prior to sampling and randomized into various timepoints between 0-45 minutes post-room disturbance to create a range of β-hydroxybutyrate levels. We collected two blood samples per individual (~140uL total) via the brachial vein using a 26-guage needle. A drop of whole blood was used to measure circulating β-hydroxybutyrate levels in duplicate for all individuals. Blood samples were subsequently centrifuged, and plasma was collected to measure β-hydroxybutyrate levels using a commercial kit (Sigma Aldrich) manufacture instructions. Due to sampling error (*e.g.,* hematocrit tube breaking), we had a total of 17 data points in which β-hydroxybutyrate levels were quantified both with the point-of-care device and benchtop assay.

Figure S1. Linear regression between benchtop assay and point-of-care measurement of β-hydroxybutyrate. The X-axis indicates values obtained via the commercial EIA kit, while the Y-axis indicates values obtained via the PrecisionXtra point-of-care device. The best fit line is indicated by a blue line, with 95% confidence intervals shown in gray shading.

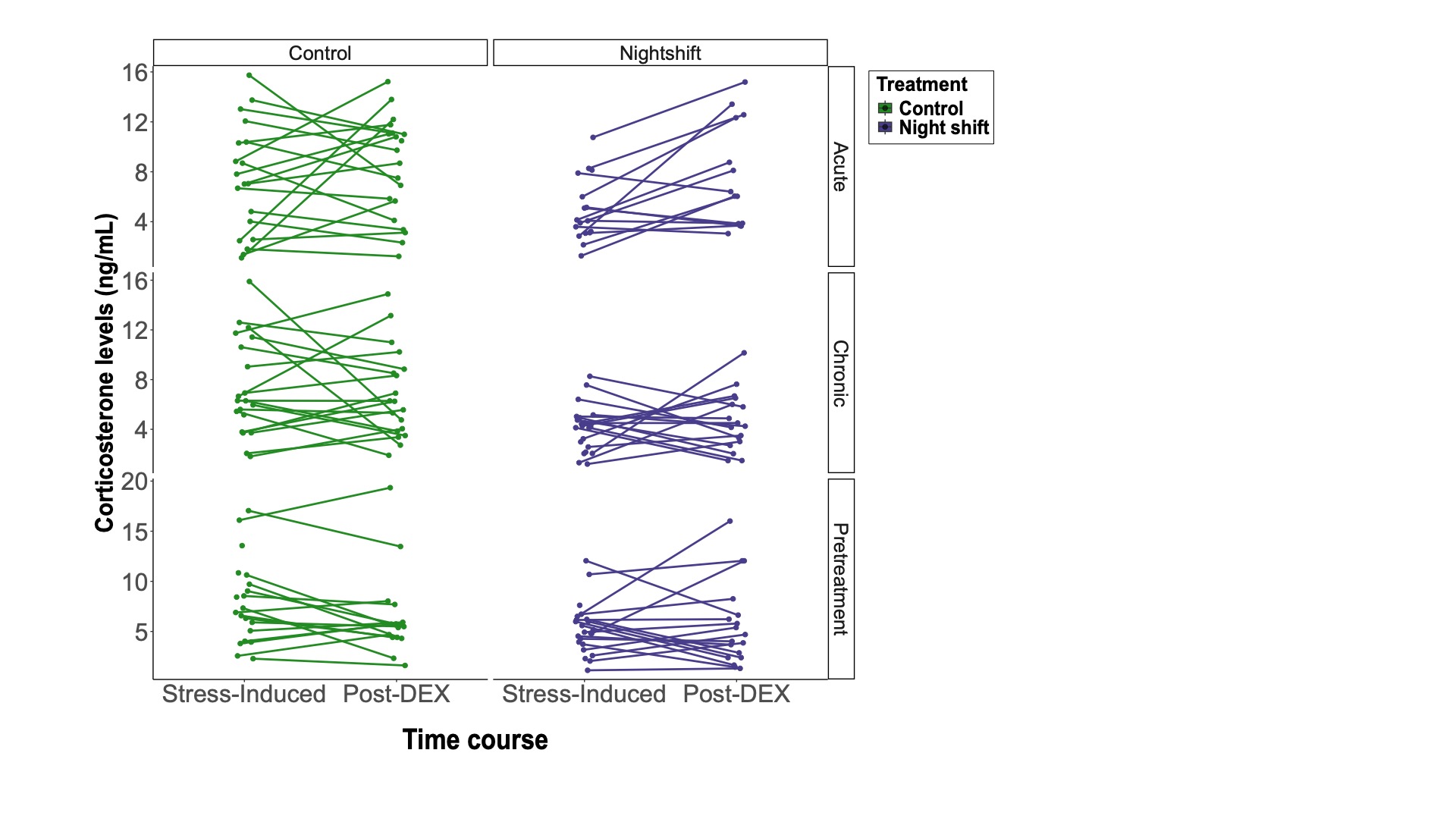

Figure S2. Raw corticosterone levels at post-restraint to after the dexamethasone challenge. The X-axis depicts the time course in corticosterone secretion, with control birds represented by a slate blue bar and night shift birds represented by a forest green bar. The Y-axis depicts corticosterone levels measured in ng/mL. Each individual point represents an individual’s corticosterone level between the two corticosterone quantification collection timepoints. Also represented is each timepoint of the experiment in which corticosterone was quantified.

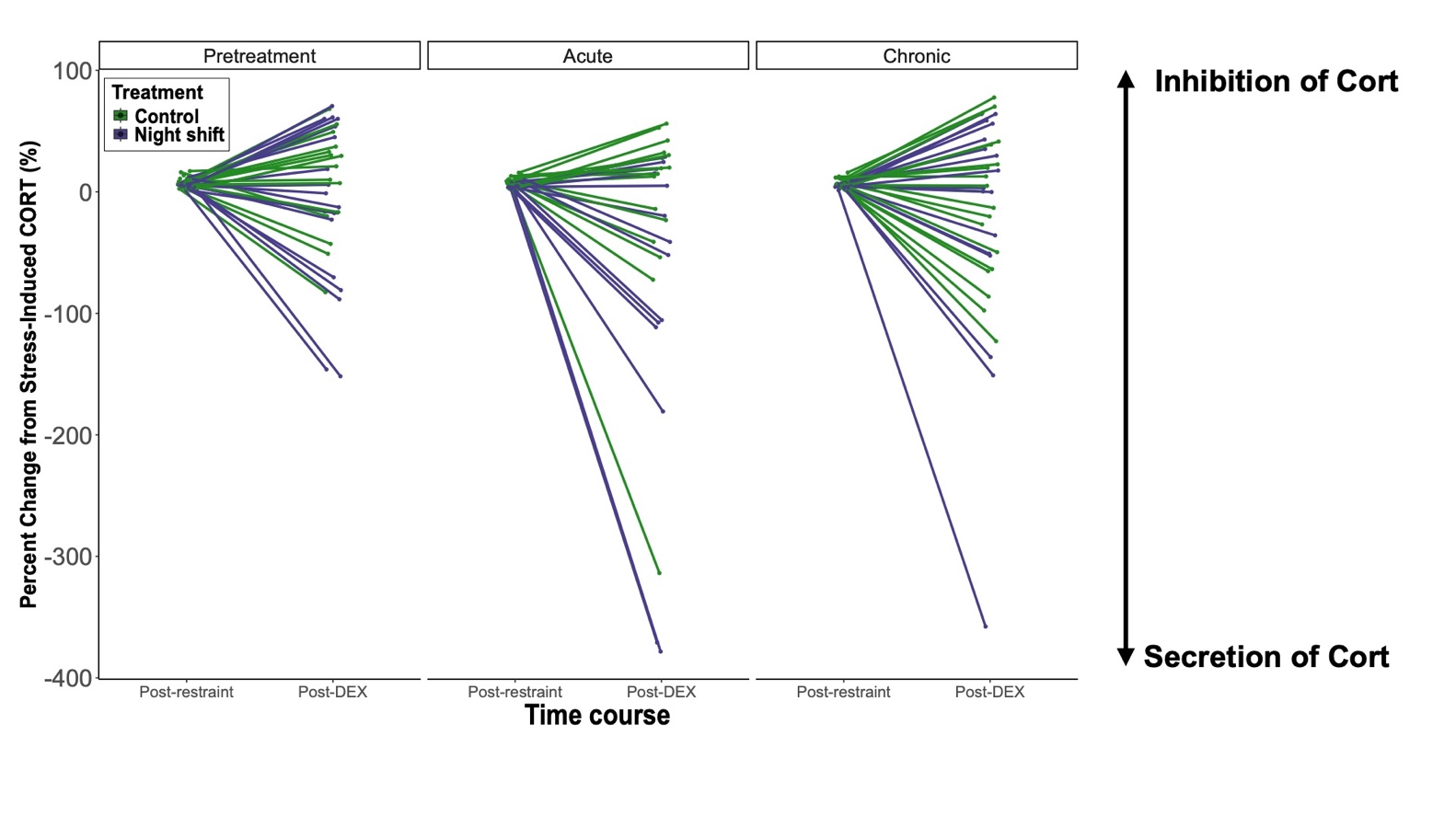

Figure S3. Relative reduction in corticosterone levels 30 min between post-restraint to after the dexamethasone challenge. The X-axis depicts the time course in corticosterone secretion, with control birds represented by a slate blue bar and night shift birds represented by a forest green bar. The Y-axis depicts corticosterone levels measured in ng/m. Each individual point represents an individual’s corticosterone level between the two corticosterone quantification timepoints. Also represented is each timepoint in which we measured corticosterone. Positive slopes represent % inhibition of corticosterone, while negative slopes indicate % increase in corticosterone secretion.

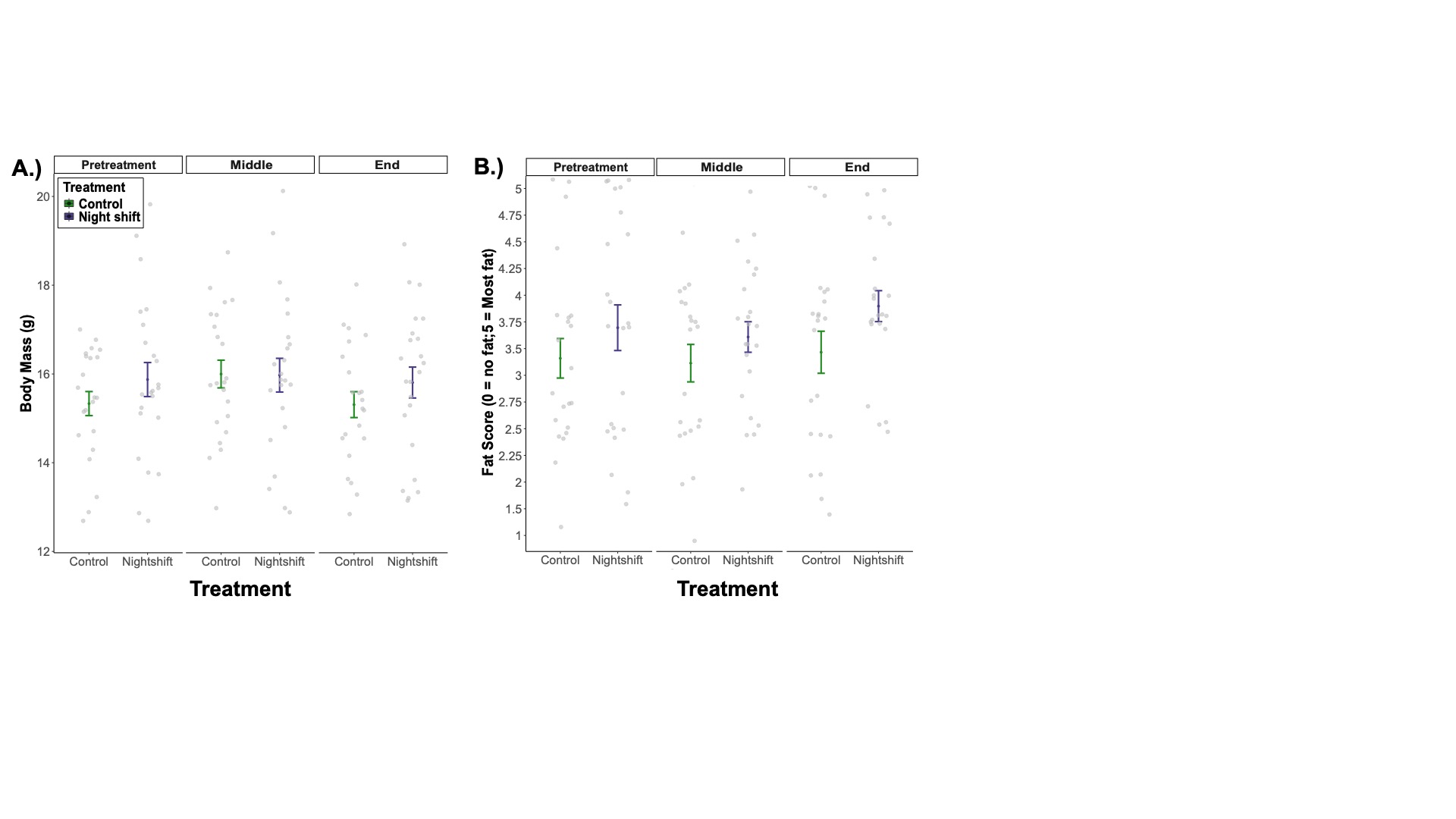

Figure S4. Morphometric measures taken at pretreatment and at death. The X-axis depicts different experimental groups, with control birds represented by a slate blue line and night shift birds represented by a forest green line. A.) The Y-axis depicts average body mass between groups at each timepoint. B.) The Y-axis depicts average fat score between groups at each timepoint. Error bars represent standard error of the mean (SEM).

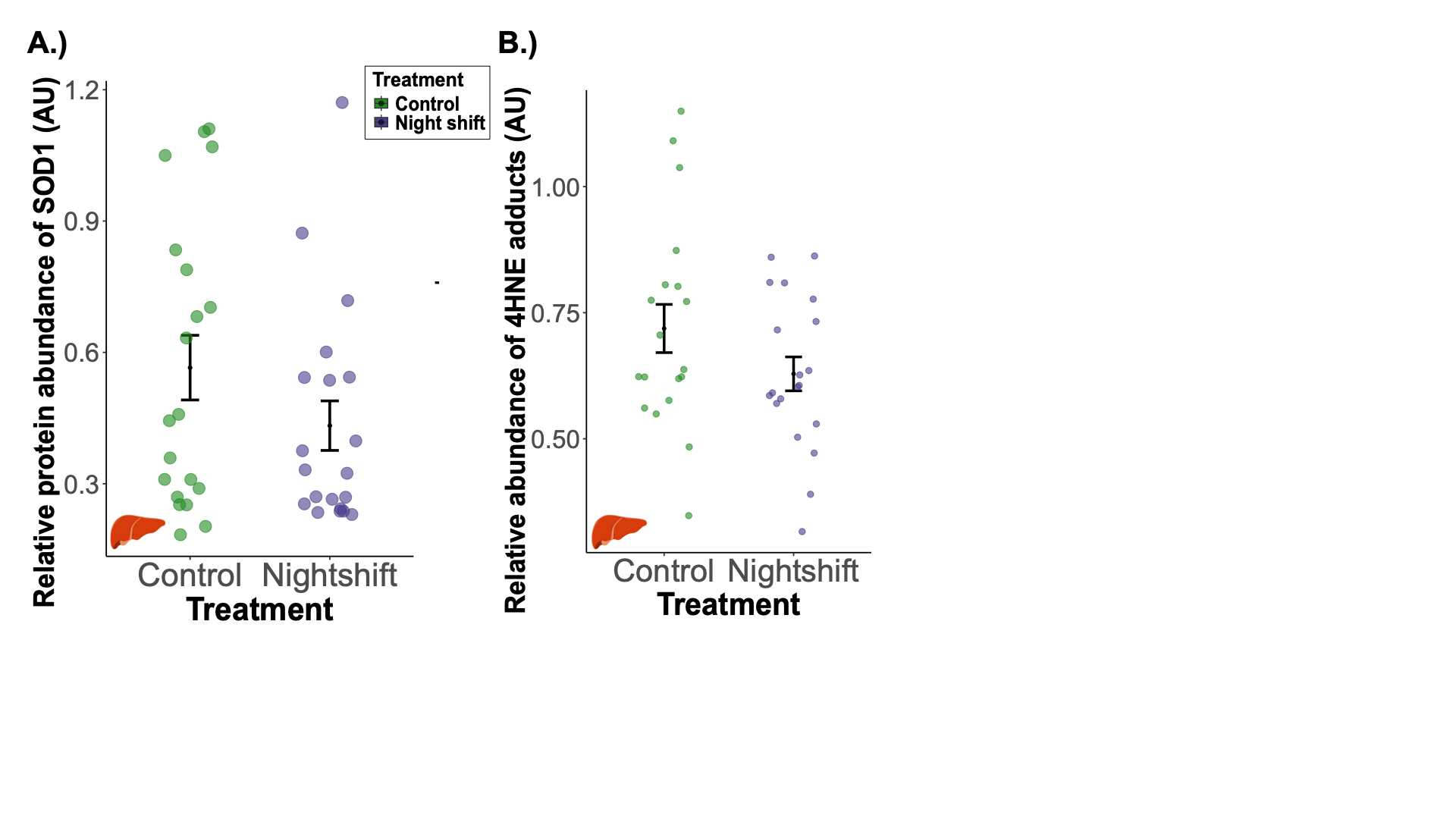

Figure S5. Relative abundance of SOD1 and 4-HNE protein levels in the liver. In both graphs, controls are depicted in dark slate blue, while birds exposed to night shift work are shown in forest green. The X-axis depicts treatment groups, while the Y-axis indicates relative protein abundance. A.) Sample size for this analysis was n = 19 for controls, and n = 20 for treatment birds. B.) Sample size for this analysis was n = 19 for controls, and n = 20 for treatment birds. Averages are plotted using raw data, with error bars indicating the standard error of the mean (SEM). Note that we show untransformed data, while in the statistical analysis, we log transform both response variables.
